# Movement Dynamics of Divisome and Penicillin-Binding Proteins (PBPs) in Cells of *Streptococcus pneumoniae*

**DOI:** 10.1101/429217

**Authors:** Amilcar J. Perez, Yann Cesbron, Sidney L. Shaw, Jesus Bazan Villicana, Ho-Ching T. Tsui, Michael J Boersma, Ziyun A. Ye, Yanina Tovpeko, Cees Dekker, Seamus Holden, Malcolm E. Winkler

## Abstract

Bacterial cell division and peptidoglycan (PG) synthesis are orchestrated by the coordinated dynamic movement of essential protein complexes. Recent studies show that bidirectional treadmilling of FtsZ filaments/bundles is tightly coupled to and limiting for both septal PG synthesis and septum closure in some bacteria, but not in others. Here we report the dynamics of FtsZ movement leading to septal and equatorial ring formation in the ovoid-shaped pathogen, *Streptococcus pneumoniae* (*Spn*). Conventional and single-molecule total internal reflection fluorescence microscopy (TIRFm) showed that nascent rings of FtsZ and its anchoring and stabilizing proteins FtsA and EzrA move out from mature septal rings coincident with MapZ rings early in cell division. This mode of continuous nascent ring movement contrasts with a failsafe streaming mechanism of FtsZ/FtsA/EzrA observed in a Δ*mapZ* mutant and another *Streptococcus* species. This analysis also provides several parameters of FtsZ treadmilling in nascent and mature rings, including treadmilling velocity in wild-type cells and *ftsZ*(GTPase) mutants, lifetimes of FtsZ subunits in filaments and of entire FtsZ filaments/bundles, and the processivity length of treadmilling of FtsZ filament/bundles. In addition, we delineated the motion of the septal PBP2x transpeptidase and its FtsW glycosyl transferase binding partner relative to FtsZ treadmilling in *Spn* cells. Five lines of evidence support the conclusion that movement of the bPBP2x:FtsW complex in septa depends on PG synthesis and not on FtsZ treadmilling. Together, these results support a model in which FtsZ dynamics and associations organize and distribute septal PG synthesis, but do not control its rate in *Spn*.

**Significance:** This study answers two long-standing questions about FtsZ dynamics and its relationship to septal PG synthesis in *Spn* for the first time. In previous models, FtsZ concertedly moves from midcell septa to MapZ rings that have reached the equators of daughter cells. Instead, the results presented here show that FtsZ, FtsA, and EzrA filaments/bundles move continuously out from early septa as part of MapZ rings. In addition, this study establishes that the movement of bPBP2x:FtsW complexes in septal PG synthesis depends on and likely mirrors new PG synthesis and is not correlated with the treadmilling of FtsZ filaments/bundles. These findings are consistent with a mechanism where septal FtsZ rings organize directional movement of bPBP2x:FtsW complexes dependent on PG substrate availability.

## Introduction

Cell division in most bacteria is mediated by the tubulin homolog, FtsZ, which polymerizes into dynamic filaments and bundles at the middle or toward the pole of dividing cells (1, 2). Polymerization of FtsZ filaments/bundles initiates sequential binding of a series of proteins that ultimately assemble into a controlled divisome machine for septal peptidoglycan (PG) synthesis leading to cell division (3, 4). The assembly of this machine involves the binding of FtsZ filaments to membrane-anchoring and filament-stabilizing and –bundling proteins (1). An ensemble of conserved FtsZ-ring component and regulator proteins then interact sequentially followed by a class B penicillin-binding protein (bPBP; transpeptidase (TP) of PG peptides), FtsW (glycosyl transferase (GT) that builds glycan chains), and MurJ (Lipid II substrate flippase) (5–7). The exact composition of FtsZ-ring divisomes, the mechanism of timing and triggering septal PG synthesis, and the involvement of PG remodeling by hydrolases is only partly understood and varies widely among different bacterial species (8–10).

Biochemical work demonstrates that FtsZ filaments move directionally by a treadmilling mechanism, similar to that first found for eukaryotic tubulins (11). In treadmilling, FtsZ-GTP monomers add to the growing (+) end of advancing filaments, and FtsZ-GDP, produced by FtsZ-catalyzed GTP hydrolysis, dissociate from the other disappearing (−) end of the FtsZ filament (12). The net result is directional movement of filaments, in which central FtsZ-GTP subunits are stationary. Total internal reflection microscopy (TIRFm) of moving FtsZ filaments/bundles and single-molecule (SM)-TIRFm experiments have recently established FtsZ treadmilling in the Gram-positive and –negative rod-shaped bacteria, *Bacillus subtilis* (*Bsu*) and *Escherichia coli*(*Eco*), respectively (12, 13). In these studies, the velocity of filaments/bundles was shown to depend on the FtsZ GTPase activity, but was independent of the addition of TP-inhibiting antibiotics. Both studies also concluded that bidirectional treadmilling of FtsZ filaments/bundles plays a role in organizing and distributing the septal PG synthesis apparatus. In *Bsu*, treadmilling is tightly coupled to and limiting for septal PG synthesis and septum closure, such that the velocity of septal bPBP2b movement correlates with the velocity of treadmilling of FtsZ filaments/bundles (12). This mode of PBP movement differs from that of MreB-mediated side-wall elongation that depends on PG synthesis and is blocked by antibiotics in *Bsu* and other rod-shaped bacteria (14, 15). Likewise, the velocities of bPBP3 (FtsI) and FtsZ treadmilling are correlated in *Eco*, but curiously, treadmilling velocity does not limit the rate of septal PG synthesis determined by incorporation of fluorescent D-amino acids (FDAAs) or the rate of septum closure (13). In contrast, after septal PG synthesis is initiated in *Staphylococcus aureus* (*Sau*), cytokinesis to close the septum does not depend on FtsZ treadmilling and is likely driven by PG synthesis (16).

Compared to these model rod-shaped and spherical bacteria, much less is known about FtsZ ring dynamics in ovoid-shaped bacteria, such as the human respiratory pathogen, *Streptococcus pneumoniae*(pneumococcus; *Spn*). Newly divided ovococcus bacteria form prolate ellipsoid-shaped cells containing equatorial rings composed of FtsZ and other proteins (bottom, Fig. S1A) (17, 18). These equatorial rings become the mature septa at the start of division (bottom, Fig. S1A) (19, 20). Mature FtsZ rings contain all of the proteins required for the stabilization and placement of FtsZ protofilaments and for PG synthesis during the next round of division (21). *Spn* lacks conventional nucleoid occlusion mechanisms, and high-resolution microscopy shows that FtsZ protofilaments are distributed in nodal patterns around mature septal FtsZ rings that surround the undivided nucleoid marked by its origin of replication (bottom, Fig. S1A) (22–24).

To construct an ellipsoid shape, two modes of PG synthesis are organized by the septal FtsZ rings in *Spn*(25). Septal PG synthesis mediated by Class b PBP2x (bPBP2x) and other proteins closes inward to separate cells, whereas peripheral PG synthesis mediated by bPBP2b and other proteins emanates outward from midcells to elongate cells (top, Fig. S1A). Early in division, a ring composed of MapZ (LocZ) splits (Fig. S1B, top; Fig. 1A) and is moved by peripheral PG synthesis toward the equators of the daughter cells (26, 27), preceded by the origin of replication (top, Fig. S1A) (23). MapZ movement precedes migration of FtsZ, FtsA (FtsZ membrane anchor and peripheral PG regulator in *Spn* (20)), and EzrA (FtsZ assembly modulator in *Bsu*(28) and FtsZ assembly positive regulator in *Spn* (A. Perez, in preparation)) to the equators (Fig. 1B-1F). During middle to-late cell division, FtsZ, EzrA, and FtsA are observed at the closing septum as well as at both developing equators, resulting in a distinctive three-band pattern (Fig. S1A, middle; Fig. S1B bottom three panels, Fig. 1B-F). After FtsZ, EzrA, and FtsA relocate to equators, proteins involved in PG synthesis, including DivIVA (negative-curvature binding protein that determines cell shape) (29), MltG (endolytic transglycosylase in peripheral PG synthesis) (30), GpsB (regulator that distributes septal and peripheral PG synthesis) (31), StkP (Ser/Thr protein kinase that regulates PG synthesis) (32), and bPBP2x (19), remain at the septum, and migrate to equators right before cells divide (Fig. 1G-L).

**Fig. 1.**
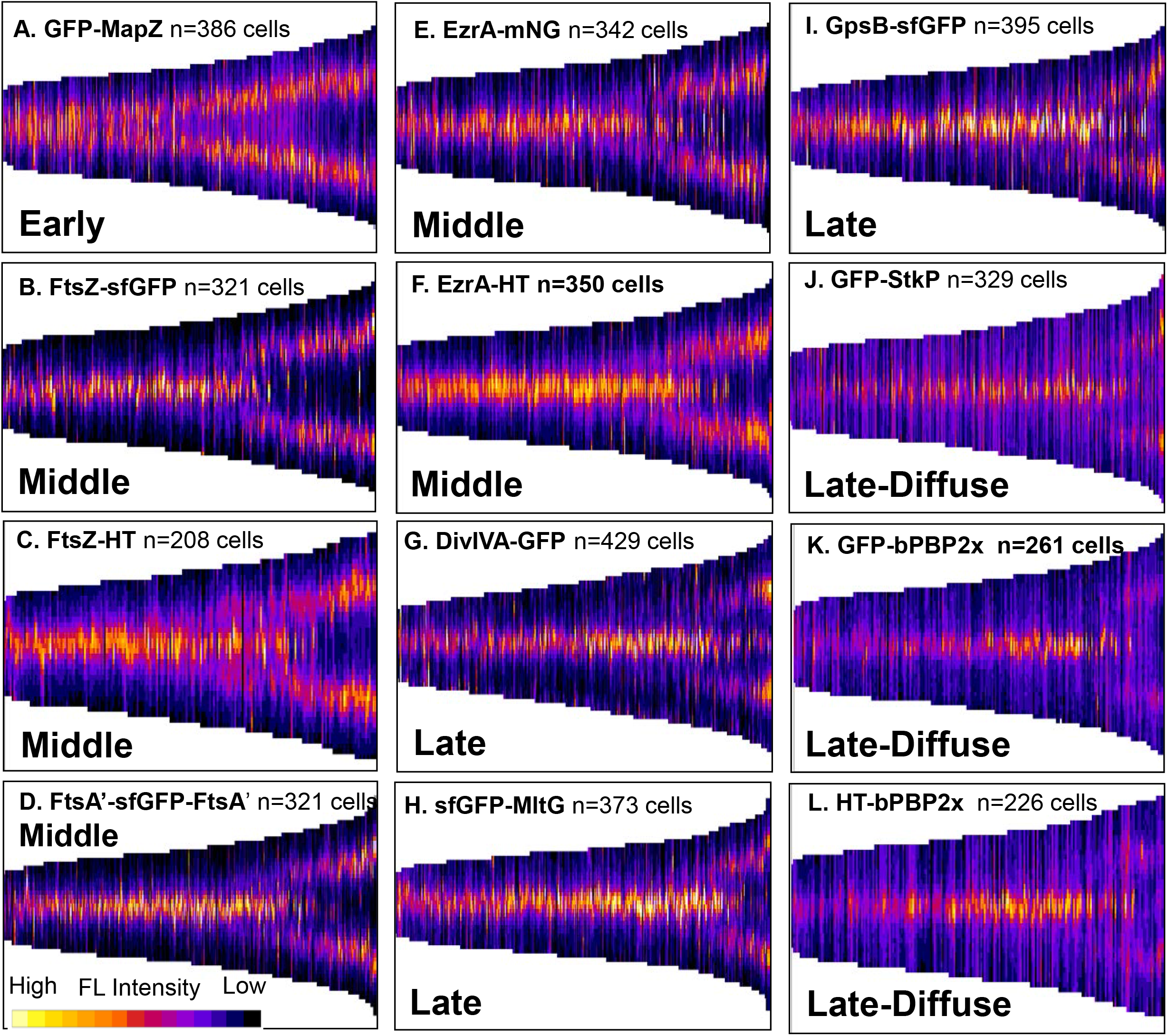
Relocation of *Spn* cell division and PG synthesis proteins occurs in three stages. Strains expressing functional fluorescent- or HT-protein fusions expressed from native loci (see Table S1; note that linkers have been omitted here to simplify nomenclature) were grown in C+Y, pH 6.9 at 37° C in 5% CO_2_ to OD_620_= 0.1–0.2, imaged, and processed by MicrobeJ to generate demographs as described in *SI Experimental Procedures*. Cells are sorted by length from shorter (left) to longer (right), corresponding to pre-divisional single cells to late-divisional daughter cells about to separate, respectively. Data are from two independent biological replicates, where the total number of cells (n) is indicated. Early arriving protein: (A) MapZ (IU9182; GFPMapZ). Middle arriving proteins: (B) FtsZ (IU9985; FtsZ-sfGFP); (C) FtsZ (IU14288; FtsZ-HT); (D) FtsA (IU13662; FtsA’-sfGFP-FtsA’); (E) EzrA (IU14117; EzrA-mNG); (F) EzrA (IU14404; EzrA-HT). Late arriving proteins: (G) DivIVA (IU9167; DivIVA-GFP), (H) MltG (IU11005; sfGFP-MltG), (I) GpsB (IU11638; GpsB-sfGFP), (J) StkP (IU9164; GFPStkP), (K) bPBP2x (IU9020; GFP-bPBP2x), and (L) bPBP2x (IU13910; HT-bPBP2x). All HT strains were labelled with HT-TMR ligand (see *SI Experimental Procedures*).

Little is known about how FtsZ moves from the septum to the MapZ rings that have arrived near the equators of daughter cells. Current models postulate that FtsZ migrates *en masse* from the septum to the equatorial MapZ rings at a later stage in division (e.g., (23)). A recent study used TIRFm to demonstrate treadmilling of FtsZ filaments/bundles in equatorial rings of *Streptococcus mutans* (*Smu*) (33), which is evolutionarily distant from *Spn*(33). In this study, *en masse* streaming of FtsZ from septa to equatorial rings was detected in a minority (≈7%) of dividing *S. mutans* cells (33). Here, we report that key proteins involved in FtsZ ring assembly and in septal and peripheral PG synthesis have different dynamics during pneumococcal cell division. We demonstrate and describe several parameters of FtsZ treadmilling in *Spn*. Furthermore, we show for the first time that nascent rings containing FtsZ, FtsA, and EzrA move out from mature septa guided by MapZ throughout the cell cycle. Streaming of EzrA filaments was only observed in Δ*mapZ* mutants as a possible division failsafe mechanism. In contrast, several other proteins were confined to mature septa and showed little dynamic movement within the limits of conventional TIRFm. Finally, we show that bPBP2x interacts with FtsW and that both proteins show directional movement along mature septal rings, independent of FtsZ treadmilling. Together, these findings reveal new aspects about the movement and assembly of FtsZ/FtsA/EzrA filament/bundles in dividing *Spn* cells and show that septal bPBP2x:FtsW complexes require PG synthesis for movement.

## Results

**Relocation of Spn cell division and PG synthesis proteins occurs in three stages and is dependent on pH**. To compare the dynamics of pneumococcal cell division and PG synthesis proteins, we constructed and vetted a large set of fluorescent- and HaloTag (HT) protein fusions expressed from single-copy genes at their native chromosome loci (Tables S1). Each protein fusion contains a linker region specified in Table S1, but omitted in the text and figures to simplify designations. An unencapsulated derivative (Δ*cps*) of serotype 2 strain D39 was used for these studies, because encapsulated D39 forms short chains (Fig. S1A) that make microscopy more difficult, and capsule tends to mask morphology defects of constructs (34). None of the final fluorescent- and HT-protein fusions ostensibly altered growth or cell morphology, and each showed localization of labeled proteins at septa and new equators of dividing cells grown exponentially in C+Y liquid medium, pH 6.9 (5% CO_2_) (Fig. 1 and S2), consistent with previous localization studies (see below).

Demographs generated by MicrobeJ (35) from fields of exponentially growing cells (e.g., Fig. S2) supported and extended the conclusion that *Spn* division and PG synthesis proteins relocate from the septa of single, early divisional cells (left side of demographs) to the equators of new daughter cells (right side of demographs) in three distinct stages (Fig. 1 and S1). MapZ relocates early, before FtsZ, FtsA, and EzrA (23, 26, 27). Residual MapZ remained between new equatorial rings until the migration of FtsZ and its associated proteins, FtsA and EzrA (Fig. 1A-1F), but a third sepal ring of MapZ reported previously (26) was not detected in cells grown under these conditions (see also (23)). FtsZ, FtsA, and EzrA next relocate to new equators at approximately the same time, with residual EzrA and FtsA remaining at septa when most of FtsZ has migrated (Fig. 1B-1F). Other cell division and PG synthesis proteins, including DivIVA, MltG, GpsB, StkP, bPBP2x, and FtsW, remain at septa after most FtsZ, FtsA, and EzrA has departed and move to the equators of daughter cells late in the division cycle (Fig. 1G-1L and Fig. S2G) (21, 22, 29, 30). The localization of StkP, bPBP2x, and FtsW is more diffuse away from septal and equatorial rings than that of the other proteins examined throughout the cell cycle (Fig. 1J-1L and S2G). Western blot control experiments did not detect cleavage of the GFP or HT reporter domains from GFP-StkP and HT-bPBP2x (Fig. S4). As shown later, diffusiveness in demographs corresponds to diffuse movement detected by TIRFm.

During these experiments, we unexpectedly noticed that the size and shape of wild-type Δ*cps* cells depends on pH in C+Y liquid medium. At pH ≈7.6 (5% CO_2_), which supports natural competence (36), pneumococcal cells are markedly longer and larger than at pH ≈6.9 (5% CO_2_), which is the physiological pH at the surface of epithelial cells in the human respiratory tract (Fig. S3A and S3B)(37). Underscoring the effects of higher pH, strains expressing GpsB-sfGFP or the GFP-StkP showed morphological defects characteristic of reduced GpsB or StkP function, respectively, in C+Y at pH 7.6, but not at pH 6.9 (Fig. S2 and S3C). Effects of pH on cell length and aspect ratio of wild-type cells were not observed in BHI broth (Fig. S3A), which we used in previous studies, but cannot be used here because of autofluorescence.

**Dynamics of FtsZ in nascent rings that form parallel to mature FtsZ septal rings**. TIRFm and epifluorescence microscopy showed that FtsZ filament motion was detected both inside and outside of mature septal rings in *E. coli*(13, 38) and *B. subtilis* cells (12). To determine the patterns of FtsZ movement in *Spn* cells, we performed comparable TIRFm, which limits illumination to an 100–150 nm slice and removes out-of-focus background fluorescence light (39). TIRFm of cells was performed on agarose pads containing C+Y, pH 7.1 (no CO_2_). Newly separated pneumococcal cells contain a mature midcell septal ring that appears as a prominent fluorescent band composed of multiple overlapping FtsZ filaments (Fig. 2A and 2B). FtsZ filament/bundle motion is detected by fluctuations in kymographs of TIRFm images (Fig. S5A and S5B), but it was not possible to reliably quantitate FtsZ filament/bundle velocities by TIRFm in densely packed mature septal rings (Fig. 2B and 2D; Movie S1). FtsZ filament/bundle speeds in mature septal rings were determined by wide-field imaging of vertically oriented cells as described below.

**Fig. 2.**
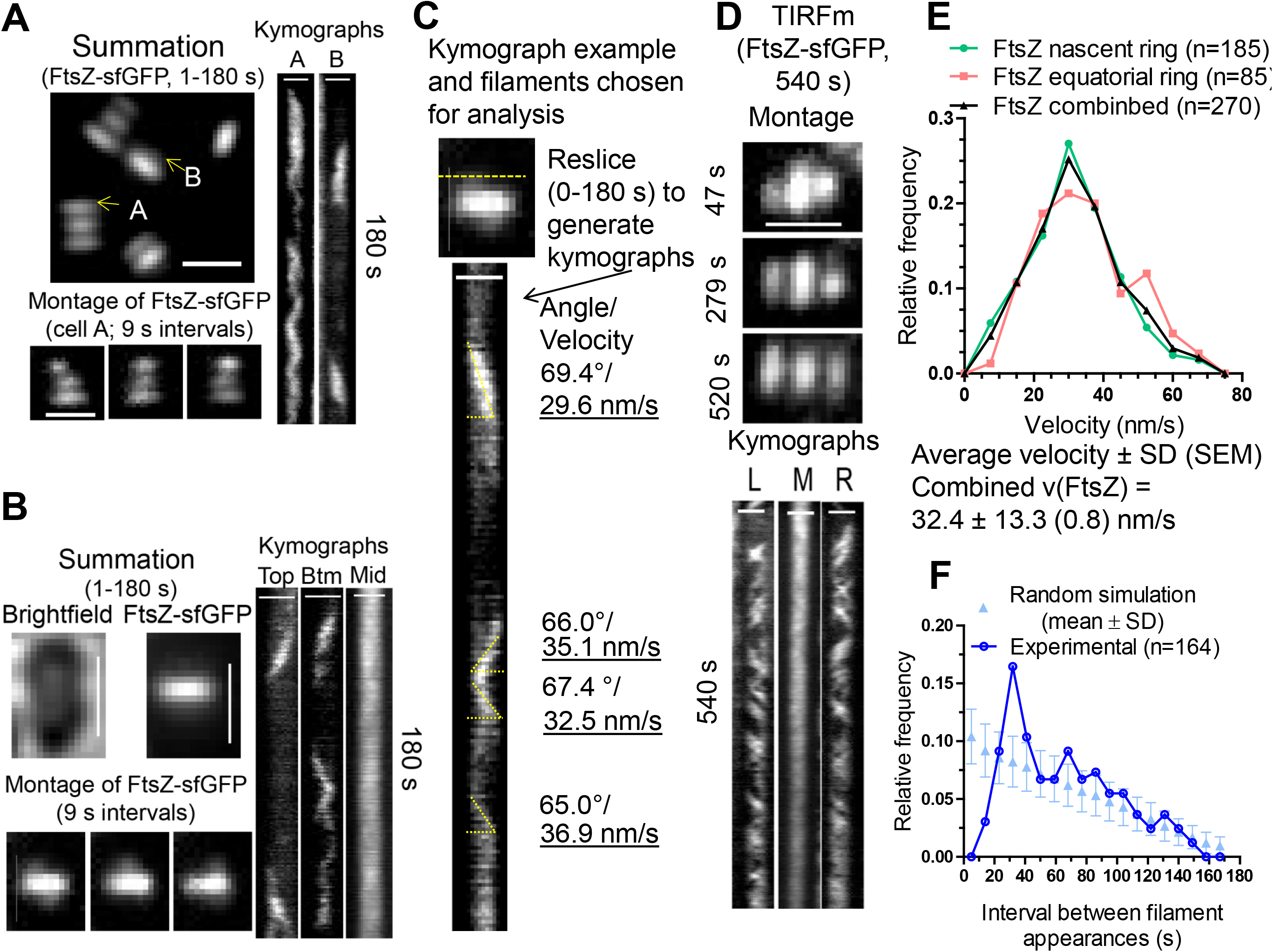
FtsZ filament dynamics in nascent and early equatorial rings determined by TIRFm of strain IU9985 expressing FtsZ-sfGFP. Representative data are shown from 2 to 4 independent biological replicates. Scale bars = 1.0 μm. (A) Summation of frames from 180 s TIRFm movies of FtsZ-sfGFP. Montage of images at 9 s intervals of the cell containing the indicated “A” ring in summation. Right panels show kymographs from 1–180 s; top equatorial “A” ring of cell in summation; area marked “B” above septal ring in summation. (B) Summation of frames from 180 s movies of bright-field images of cells and TIRFm images of FtsZ-sfGFP of mature-septal and nascent rings. Montage of images at 9 s intervals shows FtsZ-sfGFP in nascent rings above and below mature-septal ring (see Movie S1 and S2). Kymographs at right show FtsZ-sfGFP movement in the top nascent ring plane, bottom (Btm) nascent ring plane, and middle (Mid) mature-septal plane. (C) Example showing the analysis of movement of FtsZ-sfGFP filaments/bundles in a nascent ring (dotted line, top) adjacent and parallel to a mature septal FtsZ ring in an early divisional cell. Kymograph of a 180 s movie is shown. Angles were determined and converted to velocity by the equation v = tanΘ/0.079 (see *SI Experimental Procedures*). (D) TIRFm movie taken over 540 s of a single cell. Montage shows frames of FtsZ-sfGFP in two nascent rings and the middle mature-septal FtsZ ring at the times indicated. Kymographs of the left (L) and right (R) nascent rings and the middle (M) mature-septal ring over the 540 s (9 m) are shown. (E) Distribution of velocities of FtsZ filaments/bundles in nascent rings, early equatorial rings, and the combination of nascent and equatorial rings. Velocities of FtsZ filament/bundles were binned in intervals of 7.5 nm/s. (F) Distribution of times of reappearance of FtsZ filaments/bundles moving in the same direction in nascent and early equatorial rings. Reappearance times were determined as described in *SI Experimental Procedures* from four independent biological replicate experiments (n = 164 events) and are binned in 9 s intervals (dark blue). A simulation (light blue) of the means ± SDs of random events for each reappearance interval in kymographs of 1–180 s was generated as described *SI Experimental Procedures*. The reappearance interval of FtsZ filaments/bundles matched the random simulation within 2 SDs, except at 32 s, which showed a significant difference (see text).

We detected the initial stages of formation of nascent FtsZ rings on either side of mature septal rings (Fig. 2A-2D; Movie S2). FtsZ in nascent rings was detected as oblong spots moving in both directions parallel to mature septal rings (B in Fig. 2A and 2B-2D; Movie S2). Nascent FtsZ rings first appear very close to mature septal rings, and this distance increases as the nascent FtsZ filaments move outward toward the equators of daughter cells, eventually resulting in the characteristic pattern of three parallel FtsZ rings in mid-to-late divisional *Spn* cells (Fig. 2A and 2D). Summations of TIRFm images taken over 180 s movies indicate that the diameters of nascent rings start out approximately equal to those of mature septal rings (Fig. 2A and 2D). Kymographs through the long axis of cells show that nascent FtsZ rings form asynchronously on both sides of mature septal rings, with one nascent FtsZ ring detected slightly before the other ≈50% of the time (e.g., Fig. 3A). Nascent FtsZ rings move away from septal rings, add more filaments/bundles, and develop into early equatorial rings, in which directional velocities of FtsZ filaments/bundles are still detected (A in Fig. 2A; Fig. 3B). The diameters of equatorial rings became larger than those of residual septal rings, and the number of overlapping FtsZ filaments/bundles within new equatorial rings continue to increase (Fig. 2A and 3A). As the density of FtsZ filaments/bundles increases in new equatorial rings, motion is indicated by fluctuations in TIRFm kymographs (longer times, Fig. 2D; Fig. S5; Movies S1-S2). We show below that there is a correspondence between the position of nascent rings of FtsZ, FtsA, and EzrA and movement of the MapZ protein ring out from mature septal rings to the new equatorial rings of daughter cells.

**Fig. 3.**
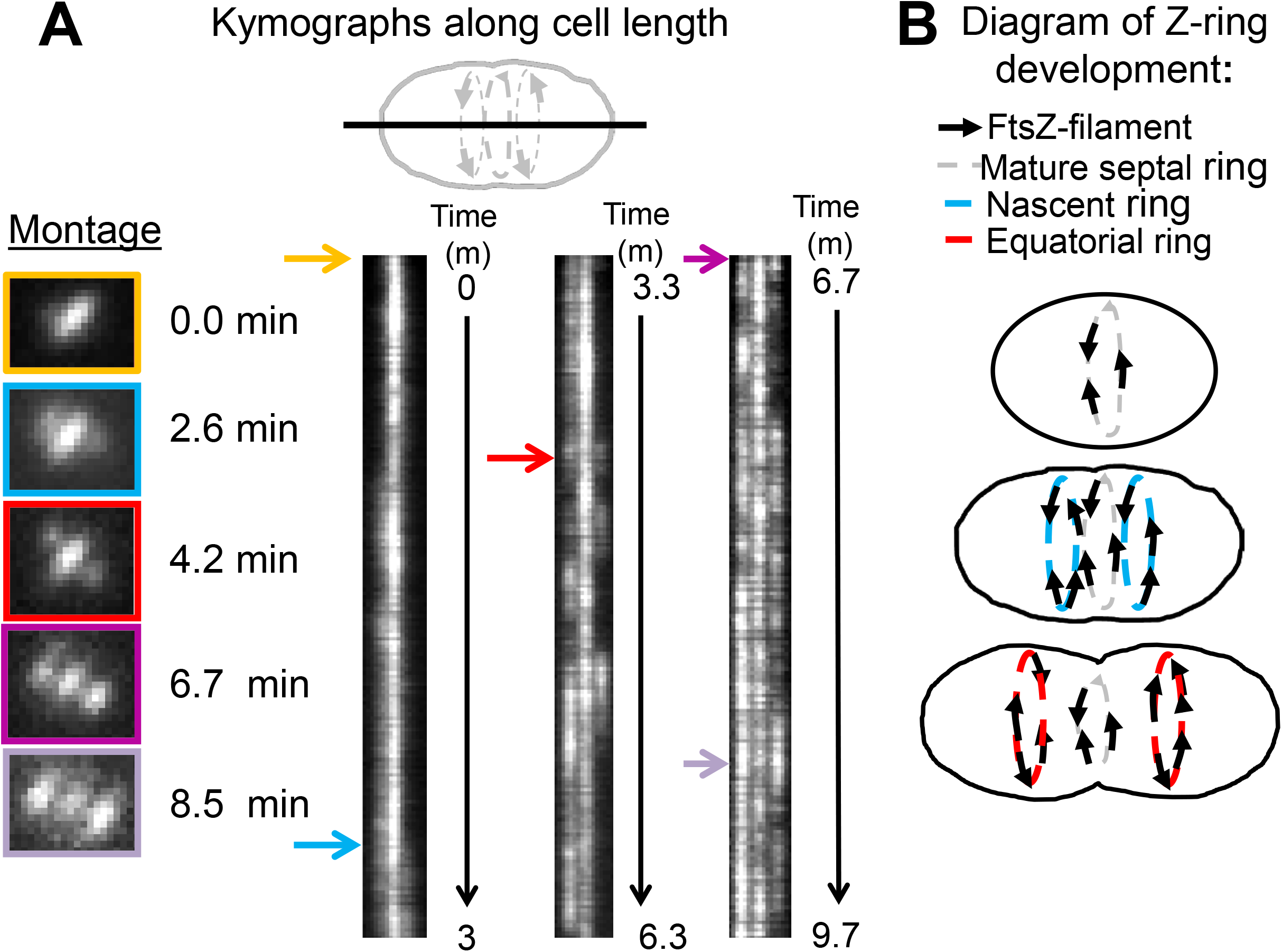
Development of nascent rings into equatorial rings of daughter cells revealed by kymographs along the cell length. TIRFm was performed to track FtsZ-sfGFP in strain IU9985 for long periods of time (9.7 m) as described in *SI Experimental Procedures*. (A) Asynchronous formation of nascent FtsZ-rings tracked by kymographs taken along the long axis of the cell. Colored arrows correspond to the times individual cells were imaged for the montage. Nascent rings formation was asynchronous in ≈50% of cells. (B) Cartoon of the continuous outward movement of nascent FtsZ rings from mature septal rings during the first stages of *Spn* cell division. See text for additional details.

Velocities of *Spn* FtsZ filaments/bundles moving in either direction in nascent rings were determined from kymographs (Fig. 2A-2D). FtsZ filament velocities were similar in nascent (31.5 ± 13.0 (1.0) nm/s (SD (SEM)) and early equatorial rings (34.4 ± 13.7 (1.5) nm/s), with a combined average FtsZ filament velocity of 32.4 ± (13.3) (0.8) nm/s in cells in C+Y, pH 7.1 (no CO_2_) (Fig. 2A-E). FtsZ filament velocities were comparable in cells in C+Y, pH 7.8 medium (no CO_2_) (33.0 ± 0.9 (10.0 nm/s); Fig. S3D). The velocities of *Spn* FtsZ filaments are similar to those reported previously for FtsZ filament/bundle movement in septal rings of *Eco* (27.8 ± 17.1 (SD)) (13) and *Bsu*(32 ± 7.8 (SD)) (12). We also analyzed the time between FtsZ filament appearances moving in the same direction (Fig. 2F). The relative frequency of appearance of FtsZ filaments moving in the same direction for the most part followed a random distribution, except between 30–40 s (Fig. 2F). The diameter of FtsZ-sfGFP rings in these live pneumococcal cells was determined by 2D-deconvolution epifluorescence microscopy to be 0.80 ± 0.06 μm, which corresponds to a circumference of ≈ 2,500 nm. Thus, the frequency of FtsZ filament appearance at intervals of 30–40 s cannot be caused by circumferential periodicity of FtsZ filaments moving at ≈34 nm/s, but may be related to an average clocked initiation of new FtsZ filaments.

**FtsZ filament/bundle dynamics and processivity in mature septal rings**. To determine the speed, processivity, and lifetime of FtsZ filaments/bundles in mature septal rings, individual *Spn* cells expressing FtsZ-sfGFP were oriented vertically in a micro-hole device described previously (Fig. 4A and S6) (12). FtsZ movement in the imaging plane was recorded by wide field time-lapse microscopy of mature septal rings with a range of diameters (Fig.4A and S7; Movie S3). Images were de-noised, and kymographs were generated (Fig. S6) (12). Lengths and angles of ≈600 FtsZ filament tracks from 29 cells were quantitated and used to compute FtsZ filament/bundle speeds, processivity, and lifetimes (Fig. 4B-4E). FtsZ filament/bundles move bi-directionally around *Spn* mature septal rings at an average speed of 30.5 ± 9.3 nm/s, which is comparable to the average velocity of FtsZ filaments/bundles in nascent and new equatorial rings (32.4 ± 13.3 nm/s; Fig. 2E) and independent of cell diameter (Fig. S7B). We conclude that the dynamic properties of FtsZ filaments/bundles in nascent and early equatorial rings match those of FtsZ filaments/bundles in mature septal rings. We also measured the total distance travelled by FtsZ filaments/bundles within septa. This gives a processivity distribution with an average of 515 ± 331 nm (Fig. 4C and 4E), meaning that an FtsZ filament typically traverses about 1/5 of the circumference of a *Spn* cell. Related to processivity, the time that FtsZ filaments/bundles exist in tracks is distributed with an average of 17.1 ± 9.4 s (Fig. 4D and 4E).

**Fig. 4.**
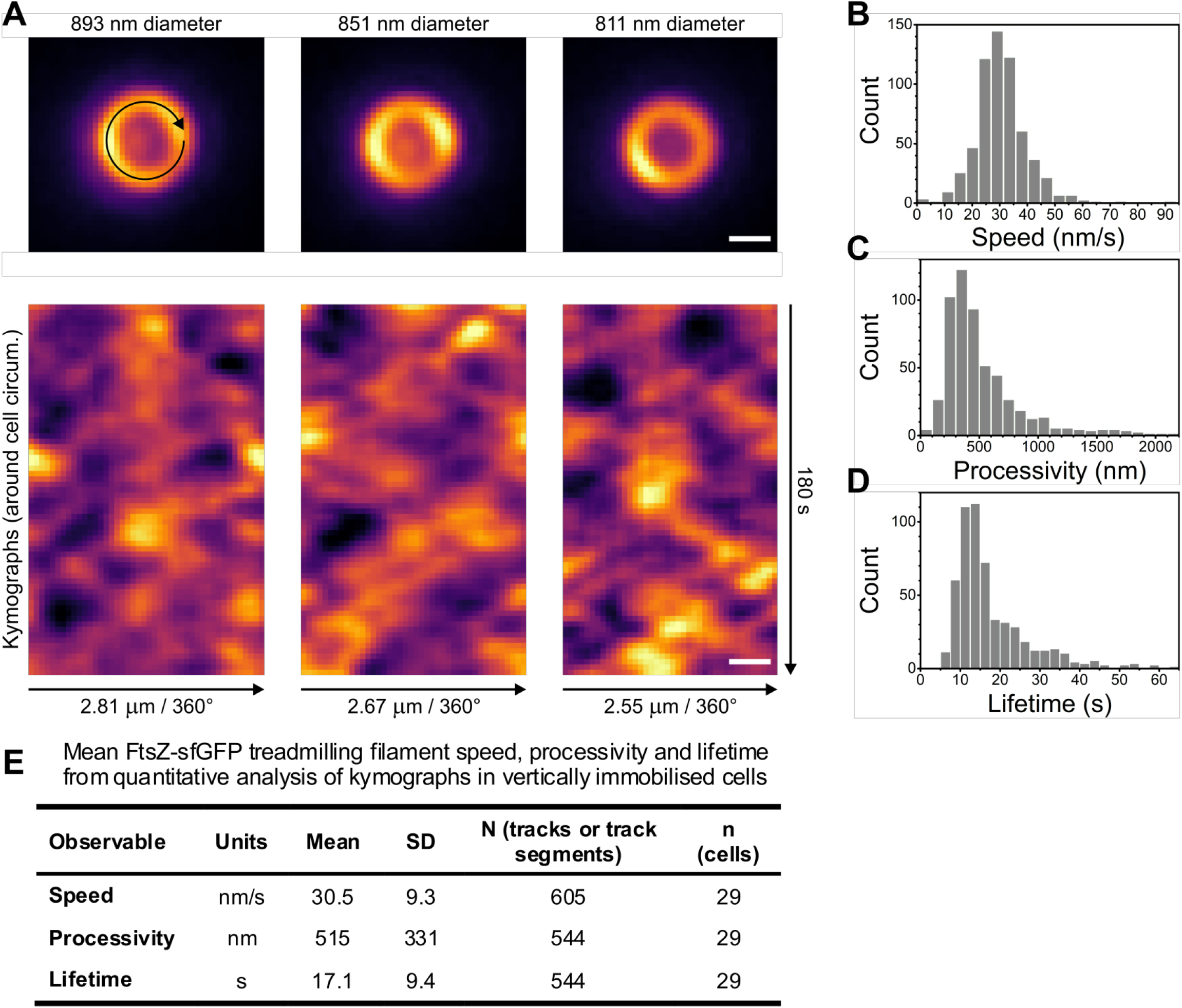
FtsZ moves bi-directionally around the mature-septal division plane with filament/bundle velocities similar to those determined in nascent FtsZ rings. IU9985 cells expressing FtsZ-sfGFP were immobilized vertically, and FtsZ-sfGFP dynamics (see Movie S3) were determined as described in *SI Experimental procedures*.(A) Top panels, representative snapshot images of FtsZ-sfGFP septal rings with typical diameters used in analyses. Images in Fig. S7 illustrate the range of cell diameters observed. Bottom panels, kymographs around cell circumference showing multiple FtsZ-sfGFP filaments treadmilling in both directions. Time lapse images of the ring circumference were unwrapped into lines (black arrow in A, top left) to generate the kymograph rows. (B-E) Individual filament tracks in kymographs were quantitated (see Fig. S6) to give FtsZ-sfGFP filament speed (B); processivity (C); and lifetimes (D), which are compiled in (E). Scale bars = 500 nm.

**Treadmilling of FtsZ filaments/bundles in mature and nascent rings and dependence of FtsZ filament velocity on GTP hydrolysis**. Previous studies show that FtsZ filaments move by a treadmilling mechanism in *Eco* and *Bsu* (12, 13). To demonstrate treadmilling of FtsZ filaments/bundles in *Spn* mature and nascent rings (Fig. 2), we constructed a functional FtsZ-HT construct (Fig. 1C) in a strain expressing EzrA-mNeonGreen (mNG) (Fig. 1E). A limiting concentration of HT substrate was added to approach single-molecule detection of FtsZ-HT by TIRFm (red, Fig. 5) in cells whose outlines were delineated by bright-field microscopy (blue, Fig. 5). EzrA-mNG was used as a fiducial marker for the locations of rings (green, Fig. 5). As presented below, FtsZ and EzrA exhibit similar patterns of movement in nascent and mature rings. In mature septal, nascent, and equatorial rings in daughter cells, single molecules of FtsZ appear as stationary foci that persist before disappearing (red spots, Fig. 5; Movie S4). We interpret these transient, static foci of single FtsZ molecules as representing non-moving FtsZ molecules within the cores of FtsZ filaments/bundles that are translocating by a treadmilling mechanism (Fig. 5B). The average lifetime of FtsZ foci detected in mature and nascent rings was 11.9 ± 9.1 (SD) s, with some foci persisting for 15–20 s (Fig. 5C). The average length of a treadmilling filament is set by the subunit lifetime and average filament speed, since subunits will bind to the plus end of a filament, and then depolymerize from the minus end. Thus, the estimated FtsZ filament length is 386 ± 335 (SD) nm.

**Fig. 5.**
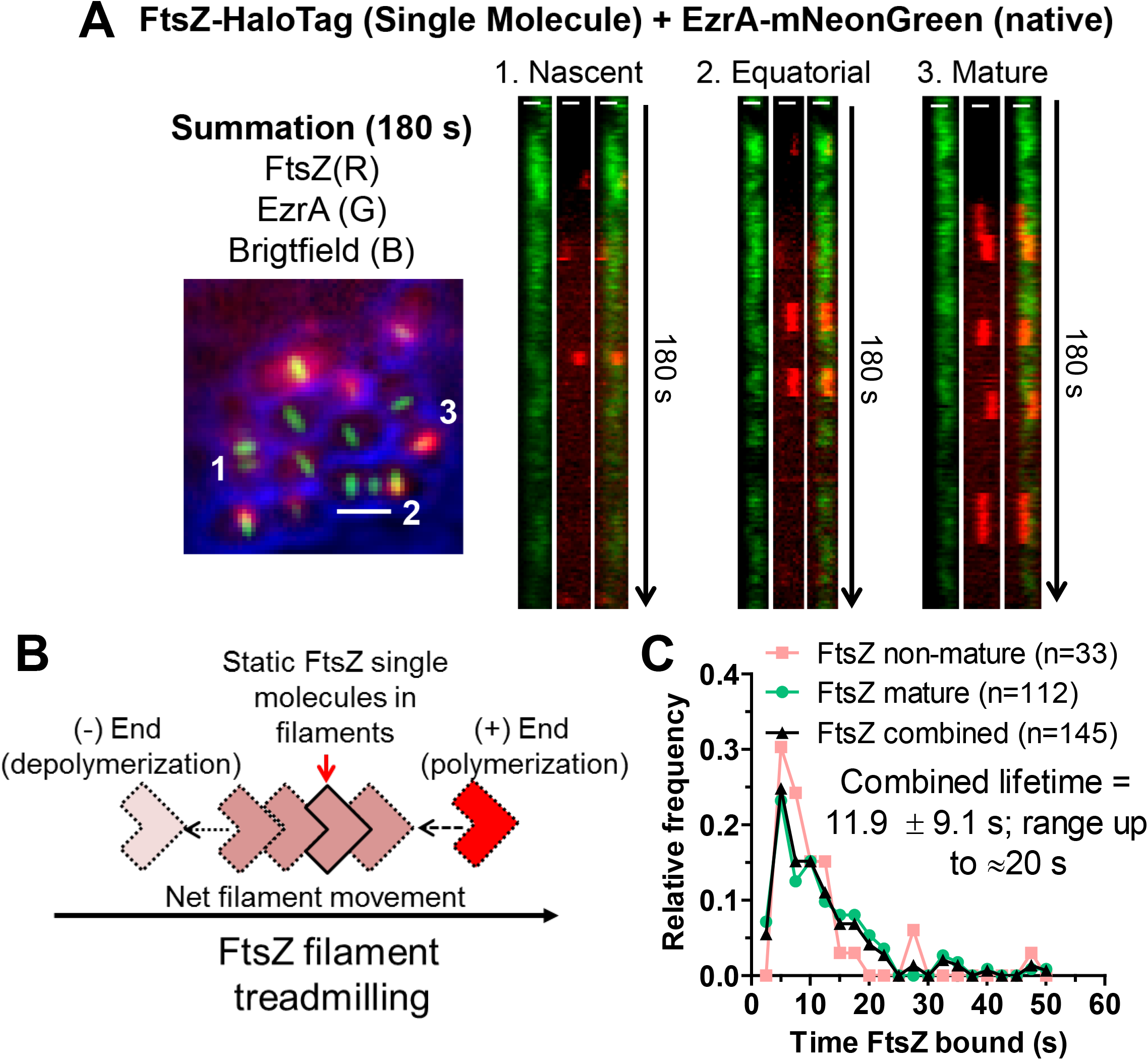
Treadmilling of FtsZ filaments/bundles in mature and nascent rings. Strain IU14352 expressing FtsZ-HT EzrA-mNG was grown in C+Y, pH 6.9 at 37° C in 5% CO_2_ to OD_620_≈ 0.1, and SM-TIRFm of FtsZ-HT labelled with JF549 HT ligand was performed on agarose pads containing C+Y, pH 7.1 at 37° C (normal atmosphere) as described in *SI Experimental Procedures*. Scale bars in summation = 1.0 μm. Left panel, overlaid summation of images from 180 s movie for bright field (blue), EzrA-mNG (green), and FtsZ-HT-JF549 (red). Right panels, kymographs from 1–180 s of rings indicated in the summation. Scale bars in kymographs = 0.5 μm. Data are representative from 2 independent biological replicates in which >60 cells were analyzed (see Movie S4). (B) Diagram of FtsZ filament movement by a treadmilling mechanism, where one molecule stays static within an FtsZ-filament that is treadmilling for roughly 12 s on average. (C) Histogram displaying the time an FtsZ single molecule is bound within non-mature rings, mature rings, and combined. Values are binned in 2.5 s increments. Labelled FtsZ-HT molecules that were present for 3 s or longer (at least 3 consecutive frames) were included in this analysis.

In addition, we confirmed that the velocity of *Spn* FtsZ filament/bundle movement depends on GTP hydrolysis by FtsZ, as reported previously for other bacteria and in biochemical reactions (12, 13, 40). For these experiments, we constructed a *Spn* mutant expressing FtsZ(G107S), which likely is defective in GTP binding based on homologues in other bacteria (Fig. S8A) (41). The *ftsZ*(G107S) mutant is temperature sensitive for growth and lyses at 42° C (Fig. S8B). Following a shift from 32° C to 42° C, the *ftsZ*(G107S) mutant formed larger, more spherical cells than the *ftsZ*^+^ parent strain (Fig. S5C and S5D), although the relative cellular amount of FtsZ(G107S) was comparable to that of FtsZ^+^ in cells at 42° C (Fig. S5E). Strains expressing FtsZ(G107S)-sfGFP are not viable. Therefore, we constructed a *ftsZ*(G107S)//*bgaA*::P_Zn_-*ftsZ-sfgfp* merodiploid strain (Table S1), in which we expressed and tracked the movement of low levels of ectopically expressed FtsZ-sfGFP by adding limited concentrations (0.1/0.01 mM) of Zn^2+^/Mn^2+^ at the still-permissive temperature of 37° C (Fig. S9). Under these conditions, the *ftsZ^+^*//P_Zn_-*ftsZ*-*sfgfp* and *ftsZ*(G107S)//P_Zn-_*ftsZ*-*sfgfp* strains show overall similar growth (Fig. S9A) and FtsZ-sfGFP localization (Fig. S9B), although slightly aberrant cells with mislocalized FtsZ-sfGFP were occasionally observed for the *ftsZ*(G107S)//P_Zn_-*ftsZ*-*sfgfp* strain. TIRFm of nascent FtsZ rings revealed that FtsZ-sfGFP filaments still move bidirectionally, but with significantly reduced velocity in the *ftsZ*(G107S) mutant compared to the *ftsZ*^+^ parent strain (Fig. S10; Movie S5). Overexpression of another mutant allele, FtsZ(D214A) that is defective in GTPase activity, also severely decreases FtsZ filament/bundle velocity (see below; Fig. S16A). We conclude that *Spn* FtsZ filament/bundle velocity produced by treadmilling is dependent on GTP binding and hydrolysis by FtsZ, consistent with previous studies in other bacteria (12, 13, 16).

**FtsA and EzrA form nascent rings with FtsZ in *Spn***. We examined the movement of several proteins involved in FtsZ filament formation and stabilization (FtsA and EzrA) and in septal peptidoglycan synthesis and cell division (MapZ, GpsB, MltG, DivIVA, StkP, bPBP2x, and FtsW) (see *Introduction*). Of this set, only FtsA and EzrA localize with FtsZ throughout the *Spn* cell cycle (Fig. 1B-1F) and form nascent rings in early divisional cells (Fig. 6; Movies S6 and S7). In summations of TIRFm movies, FtsZ, FtsA, and EzrA localize distinctly in mature septal, nascent, and equatorial rings and are not detected elsewhere in *Spn* cells (Fig. 6A and 6B). Like FtsZ, FtsA and EzrA form nascent rings moving out from mature septa (compare Fig. 2 and 6). The average velocity of EzrA in nascent rings (29.6 ± 15.3 nm/s) (Movie S6) was the same as that of FtsZ filaments (Fig. 6C). Unexpectedly, *Spn* FtsA consistently moved about 25% faster (41.7 ± 16.2 nm/s) in nascent rings than FtsZ filaments or EzrA (Fig. 6C; Movie S7), which contrasts with *Bsu* FtsA that moves at a velocity similar to FtsZ (12). We attempted to determine whether this faster velocity of FtsA is a property of the FtsA’-sfGFP-FtsA’ sandwich fusion used; however, to date, all FtsA reporter fusions constructed show growth and morphology defects. Nevertheless, we conclude that the FtsA and EzrA proteins that associate with and stabilize FtsZ filaments throughout the *Spn* cell cycle have similar overall dynamics as FtsZ filaments/bundles, including nascent ring formation (Fig. 6).

**Fig. 6.**
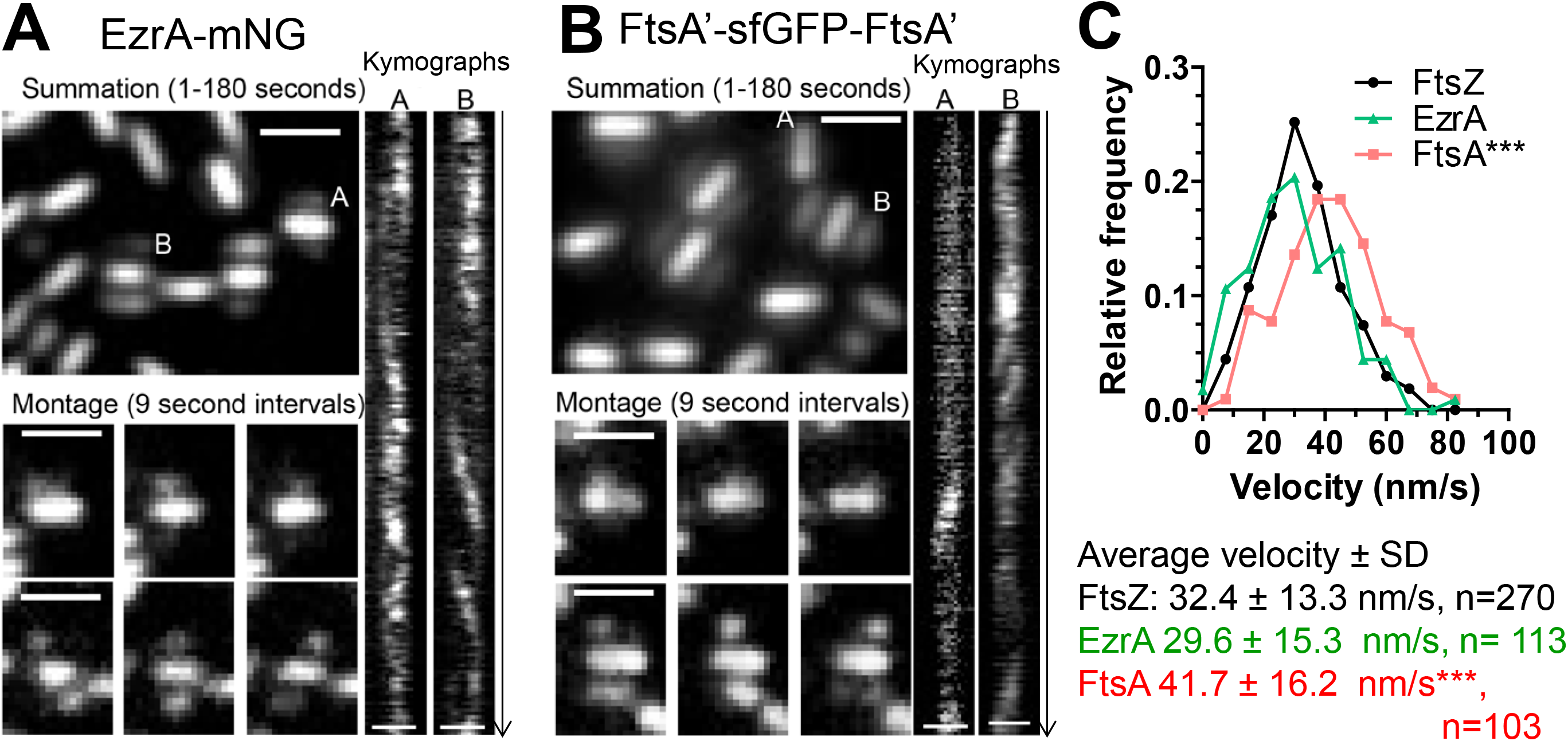
Similar movement of EzrA, FtsA, and FtsZ filaments in nascent and early equatorial rings. Movement of EzrA-mNG (IU14117) (A) or FtsA’-sfGFP-FtsA’ (IU13662) (B) were visualized by TIRFm on agarose pads containing C+Y, pH 7.1 at 37° C (normal atmosphere) as described for Figure 2. Summations are shown of individual frames of 180 s movies (see Movies S6 and S7). Montages of snapshot images show movement at 9 s intervals for nascent or early equatorial rings labeled “A” and “B” in the summations. Kymographs of the movement of EzrA-mNG or FtsA’-sfGFP-FtsA’ in these rings over 180 s are shown. Scale bars = 1.0 μm. (C) Distributions of velocities of FtsZ (n=270), EzrA (n=113), and FtsA (n=103) in nascent and early equatorial rings from four independent biological replicates. Values are binned in increments of 7.5 nm/s. P values relative to the average velocity of FtsZ-sfGFP (IU9985) (Fig. 2) were obtained by oneway ANOVA analysis (GraphPad Prism, nonparametric Kruskal-Wallis test). EzrA, no statistical difference; FtsA, P<0.001, ***.

**MapZ location corresponds to positions of nascent FtsZ and EzrA rings in early divisional *Spn* cells**. We wondered whether nascent ring formation of FtsZ, FtsA, and EzrA was coincident with movement of MapZ protein rings, which emerge from either side of mature septal rings concomitant with the start of peripheral PG synthesis (Fig. S1A) and move perpendicular to the long axis of cells to the equators of the new daughter cells (Fig. S1B and S11) (26, 27). In demographs and summations of movies, MapZ is localized primarily in mature septa or in two rings adjacent to septa, although a slight haze of MapZ remains between equatorial rings until FtsZ had fully exited from septa (Fig. 1A and S11C; Movie S8). No directional movement of MapZ or fluctuations of MapZ signal was observed in rings in kymographs (Fig. S11C and S11D), consistent with minimal MapZ movement reported previously for *Smu* MapZ (33). This conclusion was confirmed directly by SM-TIRFm of i-tag-HT-MapZ (iHT-MapZ), which unlike HT-MapZ, did not cause cell morphology defects (Fig. S11A and S11B)). Single molecules of iHT-MapZ that appeared in MapZ rings remained static for as long 60 s to >100 s before disappearing due to motion out of the TIRF plane or photobleaching (Fig. S11E and S11F; Movie S9).

In high-resolution 3D-SIM images of cells co-expressing tagged MapZ and FtsZ, low amounts of FtsZ are detected in early divisional cells at positions corresponding to nascent rings observed by TIRFm (Fig. 2; arrowhead, Fig. 7A (i) and (ii)). These nascent FtsZ rings overlap with MapZ rings moving away from septa. Likewise, EzrA in nascent rings overlaps with the parallel MapZ rings adjacent to the septum in early divisional cells (dotted box, Fig. 7A(iii)). In later divisional cells, EzrA remains at constricting septa surrounding segregating nucleoids, when all MapZ has moved to the equators of daughter cells, which also contain some EzrA (box, Fig. 7A(iv)). These results are consistent with MapZ acting as a guide for the nascent rings of FtsZ, FtsA, and EzrA that initially delivers some, but not all, of FtsZ, FtsA, and EzrA to the equators of daughter *Spn* cells.

**Fig. 7.**
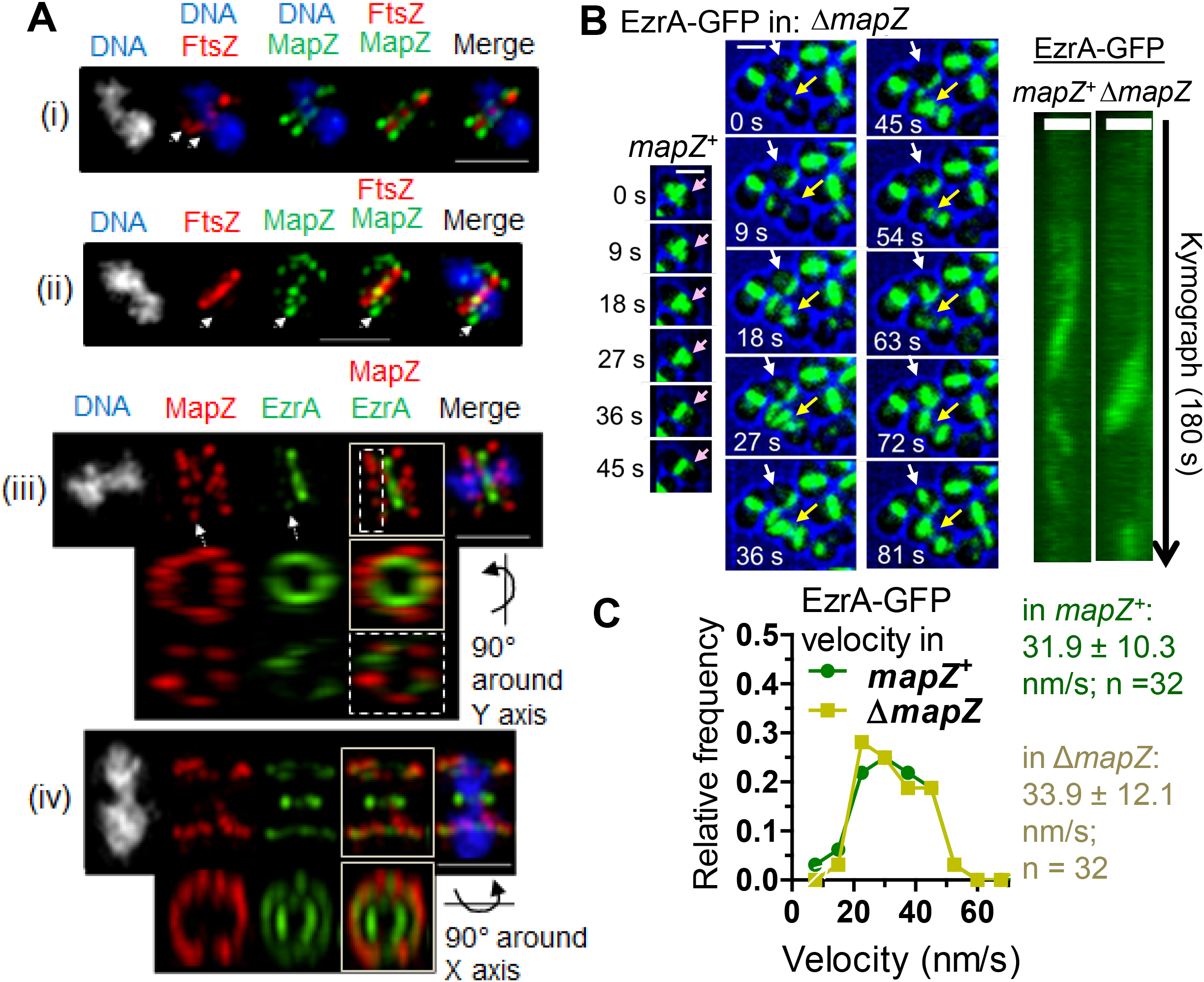
MapZ is present in nascent ring planes containing FtsZ and EzrA filaments. Immunofluorescence microscopy (A) or TIRFm (B-C) was performed to characterize the role of MapZ relative to FtsZ/EzrA filament organization. Scale bars = 1 μm.(A) Representative images of 3D-SIM dual IFM co-localizing FtsZ-Myc and MapZ-L-FLAG^3^ (strain IU9090; i and ii) or MapZ-L-FLAG^3^ and EzrA-HA (strain IU9207, iii and iv) (see Table S1). Cultures were grown in BHI, at 37° C in 5% CO_2,_ and IFM and DNA staining with DAPI were carried out as described in *SI Experimental Procedures*. For cells (i), (ii), and (iii), arrows point to the nascent ring plane where FtsZ or EzrA filaments can be seen. For cell (iii), the nascent ring (dotted box) and whole cell (solid box) were rotated, while for cell (iv), only the whole cell was rotated (solid box). The experiment was performed twice with similar results. (B) Representative montages and kymographs of EzrA-GFP (green) overlaid with bright field (blue outline, cells are black) dynamics in *mapZ*^+^ (IU10449) versus ∆*mapZ* (IU10540) strains visualized by TIRFm. Color-coded arrows highlight the location of normal EzrA-GFP filaments (pink) or cells with aberrant planes of EzrA-GFP treadmilling in streams (white or yellow arrows). Kymographs of examples of EzrA-GFP movement in *mapZ*^+^ or ∆*mapZ* cells. (C) Distributions of velocities of EzrA-GFP in *mapZ*^+^ or ∆*mapZ* strains (no significant difference; one-way ANOVA analysis, GraphPad Prism, nonparametric Kruskal-Wallis test).

If MapZ is a guide for formation of nascent rings, then we would expect aberrant movement of FtsZ/FtsA/EzrA filaments in Δ*mapZ* mutants. In the D39 *Spn* genetic background, Δ*mapZ* mutants are viable and form nearly normal looking cells with some distortions and frequent misaligned division planes (23, 42). However, FtsZ-GFP fusions in Δ*mapZ* mutants exhibit a severe synthetic defect in growth and morphology that precludes their study in *Spn* (23), but that was not commented upon in *Smu* (33). In contrast, EzrA-GFP fusions in Δ*mapZ* mutants lack this defect and appear similar to Δ*mapZ* mutants (Fig. 7B). TIRFm of EzrA-GFP movement in a Δ*mapZ* mutant indeed revealed aberrant, untimed streaming of EzrA filaments, presumably in association with FtsZ, from parent to daughter cells, often resulting in rings that are not perpendicular to the long axis of cells (arrows, Fig 7B). Nevertheless, the rate of EzrA streaming was similar in Δ*mapZ* and *mapZ*^+^ strains (Fig. 7C). Altogether, these results are consistent with MapZ acting as a continuous guide for the orderly movement of FtsZ/FtsA/EzrA filaments from mature septal rings to new equatorial rings in daughter cells. However, in the absence of MapZ, a second streaming mechanism aberrantly distributes FtsZ/FtsA/EzrA filaments into daughter cells. **bPBP2x is dynamic compared to other proteins that mediate PG synthesis**. We next examined the motion of several other proteins involved in PG synthesis in *Spn*. GpsB (regulator), DivIVA (regulator), MltG (endo-lytic transglycosylase), StkP (Ser/Thr kinase), bPBP2x (TP), and FtsW (GT) (see *Introduction*) remain at mature septa until late in the division cycle after FtsZ, FtsA, and EzrA have largely moved to the equatorial rings of daughter cells (Fig. 1G-1I; Fig. S2G). Unlike FtsZ, FtsA, and EzrA (Fig. 2, 3, and 6), TIRFm analysis did not detect GpsB, MltG, DivIVA, StkP, and bPBP2x joining nascent rings and showed that these proteins are largely confined to mature septal and equatorial rings (Fig. S12 and 8A; Movies S10-S14). In these mature septal and equatorial rings, fluctuation of GpsB or MltG signal indicative of ordered movement is not evident (Fig. S12A and S12B), whereas DivIVA and bPBP2x are actively moving, especially in equatorial rings (Fig. S12C and 8A), and bPBP2× motion is more diffusive around cells (Movie S14). By contrast, the motion of StkP is distinctively different from that of the other proteins examined. StkP locates in mature septal rings, where signal fluctuations are not readily apparent (Fig. S12D), but at the same time, StkP moves rapidly and diffusively throughout whole cells, which is captured as “clouds” of protein in summations of TIRFm movies (S12D; Movie S13). Cells expressing GFP-StkP or HT-bPBP2× did not show obvious defects in growth or cell morphology (Fig. 1 and S2), where HT-bPBP2x was labeled by an excess of HT-JF549 substrate. We conclude that of this set of proteins, only FtsZ and its ring stabilizers, FtsA and EzrA, form nascent rings and that dynamics of the other proteins varies from minimally detectable by TIRFm to diffusive. **bPBP2x and its partner FtsW move at the same velocities along septal rings**. SM-TIRFm experiments were performed to delineate the motion of bPBP2x relative to the FtsZ in mature septal rings (Fig. 8B; Movie S15). SM-TIRFm detection of HT-bPBP2× was approximated by addition of a limited amount of HT-JF549 substrate (red, Fig. 8B) that gave the same rate of bPBP2x movement when titrated downward to where labeled cells were barely detectable. FtsZ-sfGFP (green) and/or midcells determined from cell outlines (blue) were used as fiducial markers for the location of mature septal rings (Fig. 8A). Some bPBP2x molecules are detected as moving rapidly around cells in a sporadic fashion (Movie S15), consistent with TIRFm (Fig. 8B). The movement of these diffusive bPBP2x molecules was not analyzed further due to their lack of continuous tracks in SM-TIRFm. Other bPBP2x molecules attach onto mature septal rings and move directionally for at least 18 s (montage, Fig. 8B) and in some cases >30 s. The velocity of bPBP2x molecules in septal rings (21.9 ± 12.8 nm/s) is significantly slower than that of treadmilling by FtsZ filaments/bundles (32.4 ± 13.3 nm/s) (Fig. 8C). Control experiments showed that the velocity of single molecules of HT-bPBP2× is the same in strains that express FtsZ-sfGFP (Fig. 8C) or that express FtsZ^+^ (Fig. 8F).

**Fig. 8.**
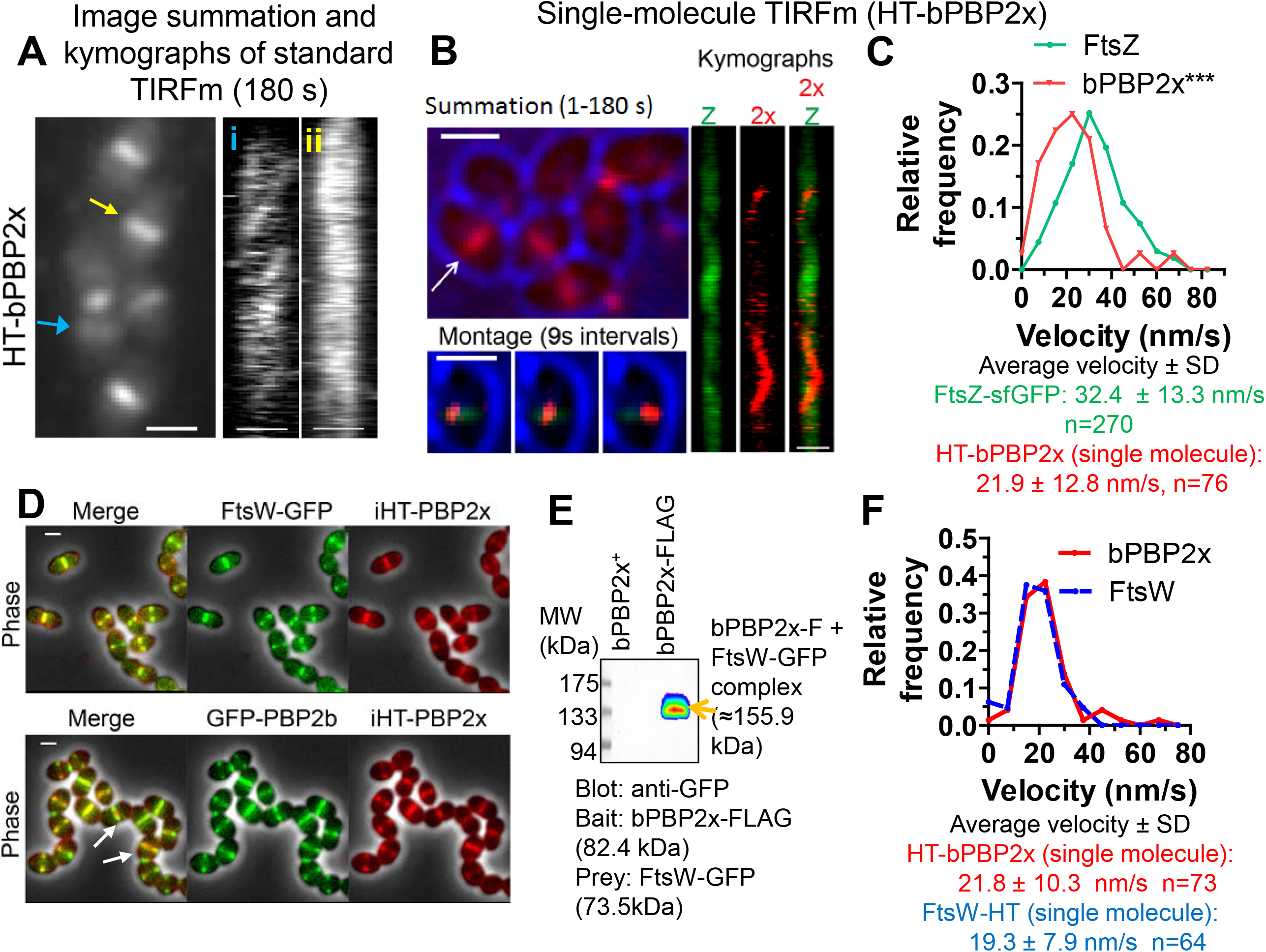
bPBP2× and FtsW septal PG synthesis enzymes co-localize, interact, and show the same movement dynamics in *Spn* cells. Microscopy experiments were performed in C+Y pH 6.9–7.1, while Co-IP was performed in BHI as described in *SI Experimental Procedures*. Scale bars = 1.0 μm. (A) Dynamic movement of multiple HTbPBP2x molecules in strain IU13910 labelled with a relatively high concentration (0.5 μM) of HT-JF549 ligand for 15 m and washed prior to imaging as described in *SI Experimental Procedures*(see Movie S14). Summation (1–180 s) with kymographs corresponding to the rings marked by colored arrows (blue/i or yellow/ii). (B) Dynamics of single molecules of HT-bPBP2x in strain IU14103 expressing FtsZ-sfGFP. Cells were labeled with a limited amount (120 pM JF549 HT ligand) and washed, and SM-TIRFm of HT-bPBP2× was performed on agarose pads containing C+Y, pH 7.1 at 37° C (normal atmosphere) as described in *SI Experimental Procedures* (see Movie S15). Summation of individual frames from 180 s TIRFm movie showing bright field (blue) outlines of cells and HT-bPBP2× (red). Montage of a series of frames at 9 s intervals corresponding to the white arrow in the summation showing a HT-bPBP2× molecule (red) moving on a mature FtsZ septal ring (green) in the middle of a cell (blue). 180 s kymographs for FtsZ (Z), bPBP2× (2×), and merged FtsZ/bPBP2x correspond to the white arrow in the summation. Scale bars = 1.0 μm. (C) Distributions of velocities of FtsZ-filaments/bundles (n=270) and single molecules of bPBP2x (n=76). Values are binned in intervals of 7.5 nm/s. FtsZ velocities are taken from Figure 2E for filaments/bundles in nascent and early equatorial rings, whereas bPBP2x velocities are taken from directionally moving single molecules in mature septal FtsZ rings, such as those shown in (B). Data are from two independent biological replicates. (D) Representative 2Depifluorescence microscopy overlaid with phase contrast images demonstrates co-localization of FtsW-GFP and itag(i)HT-bPBP2x to the inner septal ring in strain IU15066 (top row), whereas sfGFP-bPBP2b localizes adjacent to itag(i)HT-bPBP2× inner rings in strain IU15068 (bottom row). Arrows indicate iHT-bPBP2x (Red) interior to sfGFP-bPBP2b (Green). Cells were labelled with HT-TMR ligand as described in *SI Experimental Procedures*, and the experiment was performed twice with similar results. (E) FtsW-GFP (prey) is eluted with bPBP2×-FLAG (bait) in Co-IP from minimally crosslinked *Spn* cells of strain IU14964 (right lane) compared to untagged control strain (IU8918; left lane). FtsW-GFP was detected using anti-GFP (see *SI Experimental Procedures*), and the band shown was also detected with anti-FLAG antibody, confirming the presence of bPBP2x-FLAG. The experiment was performed twice with similar results. (F) Distributions of velocities of single molecules of HT-bPBP2x (strain IU13910) and FtsW-HT (strain IU15096) labelled with 120 pM of HT-JF549 ligand (see Movies S15 and S16). Values are binned in intervals of 7.5 nm/s. Data are from three to five independent biological replicates.

To further demonstrate that the velocity of HT-bPBP2x is not dependent on the fusion construct, we tracked the velocity of FtsW-HT in *Spn* (Fig. 8F; Movie S16). New results demonstrate that the biochemical GT activity of FtsW depends on its interaction with its cognate Class B PBP (43). We confirmed this interaction in *Spn* cells by co-localization of FtsW and bPBP2x (Fig. 8D) and by co-immunoprecipitation of FtsW with bPBP2× as bait in a 1:1 complex based on molecular mass (Fig. 8E). Consistent with a complex of bPBP2×:FtsW, FtsW and bPBP2× move along mature septal rings at the same velocity (Fig. 8F; Movie S16), which is slower than that of FtsZ treadmilling.

**Movement of bPBP2x and FtsW depends on PG synthesis and not FtsZ treadmilling in *Spn***. Last, we examined whether the velocity of bPBP2x and FtsW is strongly correlated with FtsZ treadmilling, as was demonstrated in *Eco* and *Bsu* (12, 13). We found that there is minimal correlation between the velocity of bPBP2x movement on septa and the rate of FtsZ treadmilling (Fig. 9A). To perform these experiments we determined the velocity of bPBP2x at septa by SM-TIRFm in FtsZ(GTPase) mutants that slowed down FtsZ treadmilling by ≈2X (FtsZ(G107S); Fig. S13 and S14; Movie S17) or ≈10X (overexpression of FtsZ(D214A); Fig. S15 and S16; Movies S18 and S19) and that lead to a percentage of cells with aberrantly placed division rings. Strikingly, reduction of FtsZ treadmilling velocity by ≈2X or ≈10X does not reduce bPBP2x velocity or reduces it only slightly (≈1.3X), respectively (Fig. 9A, S14B, and S17C). Notably, in the FtsZ(D214A) mutant, bPBP2x moves ≈5X faster than the FtsZ filaments/bundles. Likewise, FtsW velocity is only reduce by only ≈1.4X in the FtsZ(D214A) mutant (Fig. S17D). Finally, reduction of FtsZ treadmilling velocity over this range does not affect the net level of PG synthesis, as determined by incorporation of FDAA label for 2.5 m (Fig. 9B). These results contrast sharply with those for *Bsu*, where inhibition of FtsZ treadmilling significantly reduces FDAA labeling (12).

**Fig. 9.**
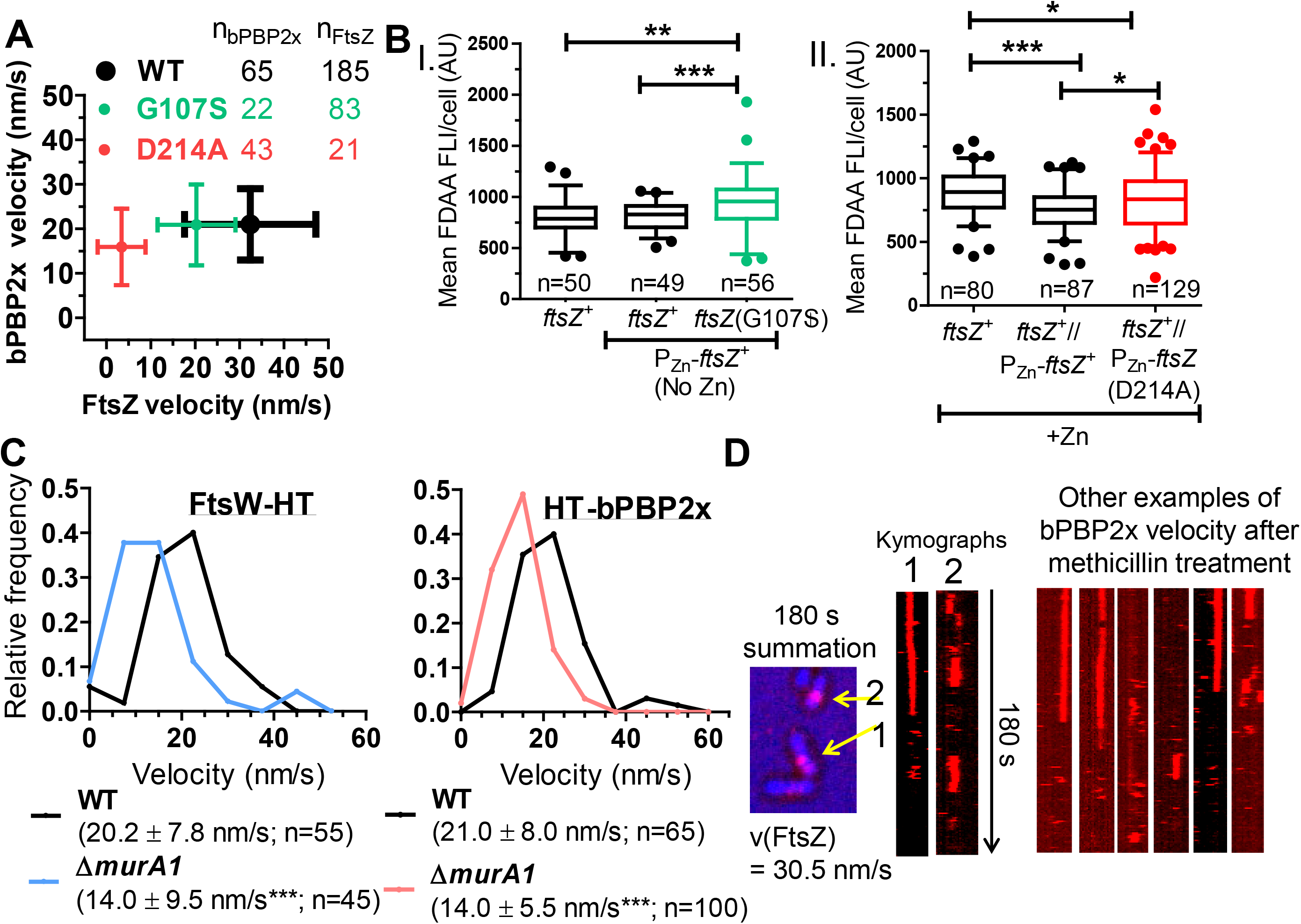
*Spn* bPBP2× and FtsW movement depend on PG synthesis and are not correlated with FtsZ treadmilling or FtsZ(GTPase) activity. Strains were grown in C+Y, pH 6.9 at 37° C in 5% CO_2_ to OD_620_ = 0.1–0.2, at which point cells were labelled with FDAA, washed, and fixed. Alternatively cells were labelled with 120 pM HT-JF549 ligand, washed, and SM-TIRFm performed to track dynamics of HT-bPBP2× or FtsWHT. (A) Mean HT-bPBP2× velocity is not correlated with FtsZ-sfGFP treadmilling velocity. Average velocities ± SDs from two or three independent biological replicates are shown for strains: IU9985 (*ftsZ*-*sfgfp*); IU13910 (*ht*-*pbp2x*); IU14375 (*ftsZ*(G107S) P_Zn_-*ftsZ*-*sfgfp*); IU14508 (*ftsZ*(G107S) *ht*-*pbp2x);* IU15181 (P_Zn_-*ftsZ*(D214A) *ftsZ-sfgfp*); and IU15041 (P_Zn_-*ftsZ*(D214A) *ht-pbp2x*) (see *SI Experimental Procedures, Table S1*, and Movies S5, S17-S19). (B) Box-and-whisker plots (whiskers, 5 and 95 percentile) of different FtsZ(GTPase) mutants showing that mean FDAA labeling of PG per cell is not reduced in FtsZ(GTPase) merodiploid mutants (middle compared to right strains). P values were obtained by one-way ANOVA analysis (GraphPad Prism, nonparametric Kruskal-Wallis test, where P>0.05, *; P>0.01, **; P>0.001, ***). Values are from two independent biological replicates (see *SI Experimental Procedures*). (C) HT-bPBP2× and FtsW-HT velocity is reduced in the absence of MurA1. Velocities were determined by SM-TIRFm in strains IU13910 (*ht*-*pbp2x*), IU15039 (∆*murA1 ht*-*pbp2x*), IU15096 (*ftsW-ht*), and IU15173 (∆*murA1 ftsW-ht*) as described inFigure 8. Shown is the average velocity ± SD of n tracks. P values were obtained by one-way unpaired, two-tailed t-tests (GraphPad Prism), where P>0.001; ***. (D) HT-bPBP2× movement is inhibited when cells are treated with methicillin. A final methicillin concentration of 0.3 μg/mL was added on top of an agarose pad after which IU13910 (*ht-pbp2×*) cells were added as described in *SI Experimental Procedures*. Cells were visualized by SM-TIRFm after 45–75 m of treatment with methicillin at 37° C (see Movie S21). A summation is shown of movie frames over 180 s with arrows pointing at septa where molecules of bPBP2x no longer move circumferentially, as indicated by the kymographs (n=57). Numbers correspond to the arrows in the summation. The experiment was performed independently twice with similar results. The velocity of FtsZ-sfGFP remained unchanged, consistent with FtsZ treadmilling, independent of PG synthesis (n=84) (see Fig S19 and Movie S20).

We next tested whether bPBP2×:FtsW velocity depends on PG synthesis. In *Spn*, there are two MurA (UDP-*N*-acetylglucosamine enolpyruvyl transferase) homologs that catalyze the first committed step of PG synthesis (44, 45). Deletion of *murA1* (*spd_0967*; also called *murZ*) does not significantly alter growth, cell morphology, or FtsZ treadmilling velocity in C+Y, pH 7.1 (Fig. S18). However, the velocity of bPBP2x and FtsW is consistently reduced by ≈1.5X in the Δ*murA1* mutant compared to the *murA1^+^* parent (Fig. 9C). We conclude that limitation of PG synthesis slows down bPBP2×:FtsW velocity without detectably affecting FtsZ filament/bundle velocity. Last, we added the β-lactam methicillin at a concentration that inhibits most of bPBP2x TP activity almost specifically (22). Methicillin addition did not inhibit FtsZ treadmilling velocity (Movie S20), but nearly completely stopped the movement of bPBP2x (Fig. 9D and S19; Movie S21). Together, these combined results indicate that movement of bPBP2×:FtsW complexes along septal rings depends on PG synthesis and is independent of the movement of FtsZ filaments/bundles.

## Discussion

Partitioning of FtsZ filaments/bundles into daughter cells occurs in ovoid-shaped (ovococcus) bacteria, such as *Spn*, by a mechanism that is fundamentally different from the Min and/or nucleoid occlusion systems present in rod-shaped bacteria (46, 47). Early in division, a ring containing MapZ protein splits from and moves parallel to the initial septal ring toward the site of the new equators in the daughter cells (Fig. S1). Movement of MapZ rings is presumably driven by outward peripheral PG synthesis from the midcell septum and is preceded by movement of the origin of replication during chromosome segregation, which is promoted by transcription and unknown mechanisms (23, 48). *Spn* MapZ is a bitopic membrane protein, whose 40 amino-terminal amino acids of its cytoplasmic domain can bind to FtsZ and whose extracellular carboxyl-terminal domain binds to PG (26, 49). However, the amino-terminal FtsZ binding amino acids are not conserved in *Smu* MapZ (33). MapZ is phosphorylated by the StkP Ser/Thr kinase; but, a requirement for MapZ phosphorylation on cell division and morphology seems to depend on *Spn* genetic background (26, 27). Likewise, in one laboratory strain, MapZ forms a third ring at division septa (26), whereas in other laboratory strains and in the progenitor D39 background of most laboratory strains, this third ring is rarely detected (Fig. 1, S7A, and S1B) (23, 27), making it unlikely that it plays an obligatory role in *Spn* division.

The interesting conjecture was made that as MapZ reaches the equators of daughter cells, it serves as a “beacon” for re-localization of FtsZ from mature septal rings (26). A recent paper proposes a concerted streaming mechanism in which FtsZ moves late in division from septa to equators in some *Smu* cells (33). In contrast, here we report that FtsZ transport to equators in *Spn* is a continuous process throughout the cell cycle (Fig. 10). Early in *Spn* division, nascent filaments/bundles of FtsZ are detected near and moving parallel to mature septal rings (Fig. 2 and 10). In the ≈10 m interval (1/3 of a generation) between the initial movement of MapZ and the migration of most FtsZ to equators (Fig. 1A and 1B), nascent FtsZ filaments/bundles move outward and become more dense until they reach equators, after which the remainder of FtsZ migrates to form mature equatorial rings (Fig. 3). Progressive nascent ring formation was detected in both Δ*cps* derivatives and in the progenitor encapsulated *cps^+^* parent D39 strain. These nascent FtsZ rings also contain EzrA and FtsA, which bind to, membrane anchor, and stabilize FtsZ filaments/bundles (Fig. 6), but none of the other PG synthesis proteins analyzed in this study was detected moving in nascent rings. FtsZ and EzrA move at the same velocity, whereas FtsA seems to move faster (Fig. 6C). Because strains expressing other FtsA fusions showed phenotypes, we could not distinguish whether this faster velocity of FtsA was construct specific. However, *Spn* FtsA has the unique biochemical property of readily polymerizing into filaments (50), and the faster velocity of FtsA could reflect this additional polymerization in cells.

**Fig. 10.**
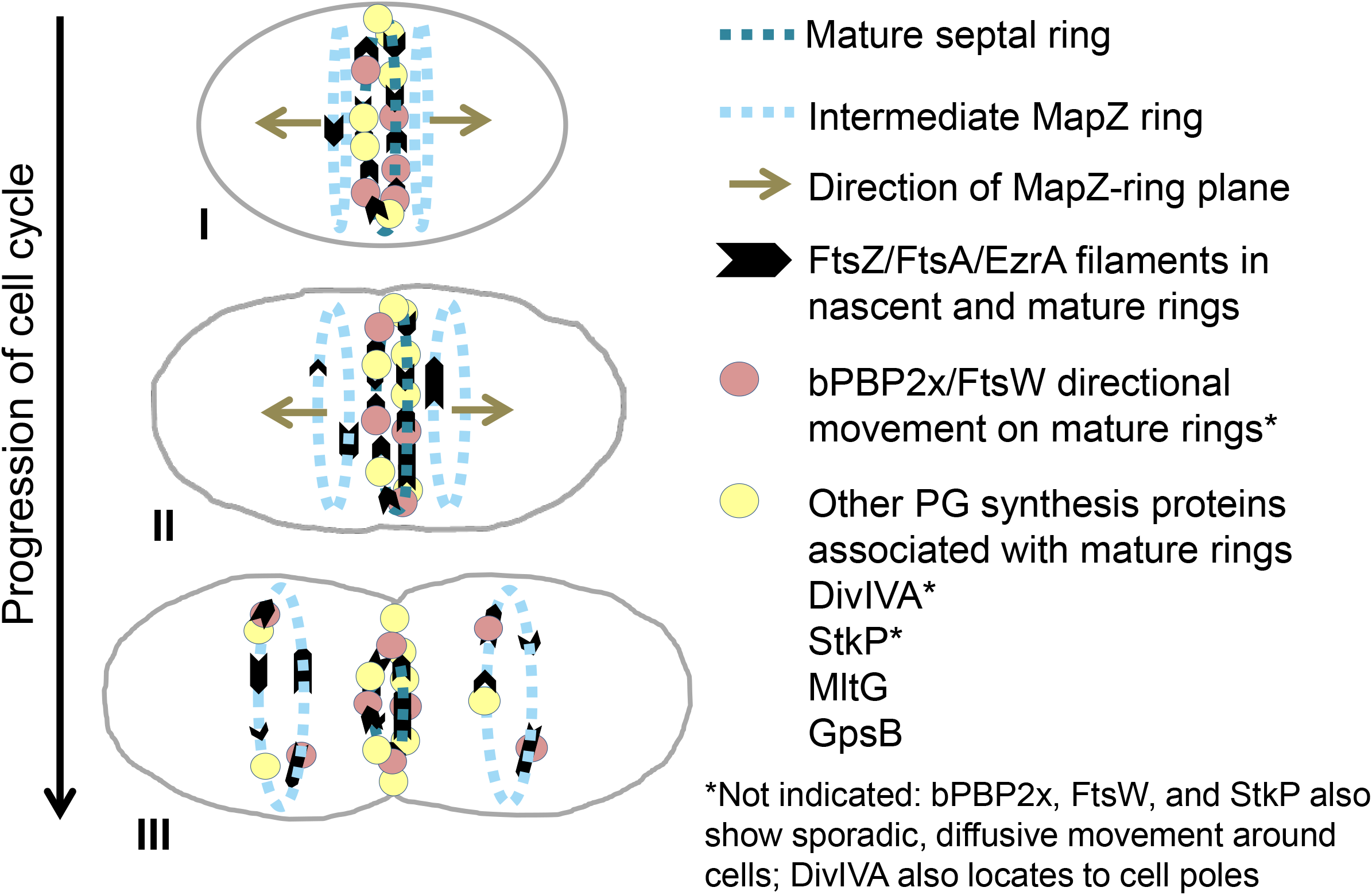
Summary diagram of the movement dynamics of FtsZ, FtsA, EzrA, bPBP2x, FtsW and regulators of PG synthesis in *Spn* cells. The division cycle is simplified to three stages. In early divisional cells (I), the equator becomes the septum of dividing cells, and most divisome proteins locate to the mature septum, with the exception of StkP and bPBP2x, which also exhibit sporadic, diffusive movement throughout cells. bPBP2x and FtsW molecules attach to the mature septal ring and move in one direction or the other for as long as 30s. After the start of peripheral PG synthesis, MapZ bifurcates into rings on both sides of the mature septal ring, and the MapZ rings start to move outward toward the positions of the equators in the daughter cells (I-II). Concurrently, nascent filaments/bundles consisting of FtsZ and its associated proteins FtsA and EzrA are detected in MapZ rings, suggesting that MapZ rings continuously guide a fraction of FtsZ/FtsA/EzrA to new equatorial rings, where FtsZ may nucleate the transport of remaining FtsZ from the mature septal ring later in division. Treadmilling velocity and dynamics of FtsZ and EzrA are the same in mature septal, nascent, and equatorial rings, whereas FtsA may move faster than FtsZ in nascent rings. FtsZ/FtsA/EzrA filaments/bundles accumulate as nascent MapZ rings move away from the mature septal ring and become early equatorial rings in daughter cells. Directional movement in or association with nascent rings was not detected for PG synthesis proteins bPBP2x, FtsW, DivIVA, MltG, StkP, and GpsB, and these PG synthesis proteins remain at constricting septa until migrating to new equatorial rings late in cell division, where bPBP2x and FtsW form a complex and move circumferentially in either direction (III). The movement of bPBP2x:FtsW complexes in septal PG synthesis depends on and reflects new PG synthesis and is not correlated with the treadmilling of FtsZ filaments/bundles. See *Discussion* for additional details.

High-resolution microscopy and effects of a Δ*mapZ* mutation suggest that nascent FtsZ/FtsA/EzrA filaments/bundles use MapZ as a guide, as opposed to a beacon, to reach the equators of daughter cells. 3D-SIM IFM images detect FtsZ and EzrA together with MapZ in early and later nascent rings (Fig. 7A). Furthermore, in the absence of MapZ, orderly nascent EzrA rings are lost, and EzrA abruptly streams between daughter cells, often resulting in the aberrant ring orientation reported previously (Fig. 7B) (23). Streaming rarely was observed in wild-type *Spn* cells (≈1%) and represents a second mechanism for translocation of FtsZ and its associated proteins to daughter cells. In this respect, streaming is a “failsafe” mechanism that accounts for the lack of lethality of *Spn* Δ*mapZ* mutations.

In *Smu* cells, continuous FtsZ nascent ring formation was not reported and streaming, which was detected in ≈7% of cells, is proposed as the primary mechanism for FtsZ movement from septal to equatorial rings (Fig. 6 in (33)). A possible reason for this difference is that *Spn* and *Smu* are evolutionarily distant *Streptococcus* species (33), and ovococcus bacteria exhibit differences in the relative timing of septal and peripheral PG synthesis (51). On the other hand, technical or strain differences may underlie the different results. In particular, the *Smu* cells in movies shown in (33) appear to be in middle-to-late divisional stages and contain prominent equatorial rings, whereas movement of *Spn* FtsZ was recorded in early to late stages of division in this study (Fig. 2 and 3). Together, our results indicate that treadmilling FtsZ/FtsA/EzrA filaments/bundles are components of migrating MapZ rings throughout the cell cycle in *Spn* (Fig. 10) and may thus, play an important role in assembly and organization of these rings, about which little is known. This transport mechanism also moves part of the cellular FtsZ population to the equatorial rings, where the FtsZ filaments/bundles may serve to nucleate the transport of the remainder of septal FtsZ later in the division cycle (Fig. 10). This movement likely occurs mainly by depolymerization of FtsZ filaments/bundles and possibly by some new protein synthesis, since concerted streaming of FtsZ filaments/bundles was rarely observed in wild-type *Spn* cells.

In the course of these studies, we also determined a set of basic parameters about FtsZ dynamics in *Spn* cells. Filaments/bundles of wild-type FtsZ always move with a velocity of ≈31 nm/s in rings at all stages of division and while streaming in Δ*mapZ* mutants (Fig. 2E, 4B, and 7C). The same velocity of FtsZ filament/bundle movement was determined by TIRFm (Fig. 2E) and independently by wide-field observation of vertically immobilized cells (Fig. 4). A treadmilling mechanism of FtsZ filament/bundle movement was confirmed directly by SM-TIRFm (Fig. 5), and SM-TIRFm and immobilized cell measurements indicated the lifetime of FtsZ subunits in filaments/bundles as ≈12 s (Fig. 5C) and the lifetime of entire FtsZ filaments/bundles as ≈17 s (Fig. 4D and 4E). Based on the average subunit lifetime and the velocity of filament/bundle movement, the average length of treadmilling FtsZ filaments/bundles in *Spn* cells is ≈390 nm. In addition, our measurements show for the first time that the processivity of treadmilling of FtsZ filaments/bundle is ≈500 nm (Fig. 4C), indicating that FtsZ filaments/bundles traverse ≈20% of the circumference of *Spn* cells on average. Finally, as expected from previous precedents (12, 13), mutations that decrease GTP binding or the GTPase activity of FtsZ reduce the velocity of FtsZ treadmilling by as much as 10X (Fig. 9A). FtsZ(GTPase) mutations also disrupt the placement of division planes compared to wild-type cells (Fig. S9, S13, and S15).

Besides FtsZ, FtsA, and EzrA, none of the PG synthesis proteins tested in this study was a member of translocating MapZ rings (Fig. 8A and S12). Within the limits of standard TIRFm, some of these proteins showed minimal movement in mature septal and equatorial rings (i.e., GpsB (regulator), MltG (endo-lytic glycosylase), and StkP (Ser/Thr kinase)) (*Introduction*; Fig. S12), whereas DivIVA (regulator) and bPBP2x (septal TP) showed obvious dynamic movements (Fig. 8A and S12C). We therefore determined the role of FtsZ treadmilling on the motion of bPBP2x and its partner FtsW in mature septa of *Spn* cells. A new biochemical study reports that FtsW GT activity depends on interactions with its cognate Class B PBP (43). In support of this interaction in *Spn* cells, bPBP2x and FtsW colocalize at all stages of *Spn* division (Fig. 8D) and bPBP2x pulls down FtsW in a likely 1:1 complex (Fig. 8E). In addition, SM-TIRFm showed that bPBP2x and FtsW move at the same velocity on septa (Fig. 8F). Moreover, besides possibly stimulating FtsW transglycosylase activity in *Spn* cells, the TP activity of bPBP2x is required for septal PG synthesis, because a *pbp2×*(S337A) active-site mutant is not viable (H.-C. Tsui, unpublished result).

Five pieces of evidence support the conclusion that movement of the bPBP2x:FtsW complex in septa of *Spn* cells depends on PG synthesis and not on FtsZ treadmilling. First, the velocity bPBP2x and FtsW is slower than that of FtsZ treadmilling in wild-type *Spn* cells (Fig. 8C). Second, the decreased velocity of FtsZ treadmilling in FtsZ(GTPase) mutants is not correlated with a decrease of bPBP2x velocity (Fig. 9A). In fact, in the slowest mutant (FtsZ(D214A) overexpression), bPBP2x is moving about 5X faster than FtsZ treadmilling. Third, severe reduction in FtsZ treadmilling velocity does not markedly decrease PG synthesis indicated by FDAA incorporation (Fig. 9B). Fourth, a decrease in PG synthesis precursors caused by a Δ*murA1* mutation, decreases the velocity of bPBP2x and FtsW by the same amount, but does not decrease FtsZ treadmilling rate (Fig. 9C). Last, addition of methicillin at a concentration that mainly inhibits bPBP2x TP activity stops the movement of bPBP2x, but does not decrease the velocity of FtsZ treadmilling (Fig. 9D).

These results strongly support the conclusion that the movement of the bPBP2x:FtsW complex in septal PG synthesis in *Spn* cells depends on and likely mirrors new PG synthesis and is not correlated with the treadmilling of FtsZ filaments/bundles. By contrast, the velocities of the septal Class B PBPs of *Bsu* and *Eco* are coupled to and limited by FtsZ treadmilling, resulting in a correlation between septal bPBP and FtsZ treadmilling velocities (12, 13). The mechanism(s) underlying this coupling and its relationship to the rate of PG synthesis in *Bsu* and *Eco* are not understood. On the one hand, FtsZ treadmilling is further coupled to and limiting for septal PG synthesis and the constriction of *Bsu* cells (12). On the other hand, the velocity of FtsZ treadmilling is not correlated with the rate of PG synthesis determined by FDAA incorporation or the rate of septum closure of *Eco* cells (13, 52). These differences suggest that additional metabolic (e.g., PG precursor pools) and structural (e.g., PG width and outer membrane synthesis) constraints may influence the relative rates of FtsZ treadmilling, bPBP complex movement, and PG synthesis in different bacteria (53, 54).

Besides the sidewall Rod complexes of rod-shaped bacteria (14), there is another precedent for the dependence of PBP movement on PG synthesis. Recent results show that septal PG synthesis continues to close division septa of *Sau* after FtsZ treadmilling is inhibited by addition of a drug (PC190723) (16). This finding is again consistent with an FtsZ treadmilling-independent mechanism by which PG synthesis itself drives for PBP motion (13, 52). The dependence of bPBP2x:FtsW movement on PG synthesis can be rationalized by a model proposed for the dependence of PBP movement on PG synthesis in sidewall elongation of rod-shaped bacteria (14). It was proposed that MreB filaments form tracks that direct the linear motion of PBP complexes (14, 55). At any point in a track, the PBP complex has used substrate behind it to synthesize PG, and the utilization of available substrate in front of it drives its motion. In the case of *Spn* bPBP2x:FtsW, it is possible that FtsZ filaments/bundles themselves or other proteins in septal FtsZ rings provide tracks that couple movement to PG synthesis. In this model, FtsZ treadmilling acts to dynamically distribute and spatially organize directional PG synthesis, possibly through indirect interactions, as suggested by recent high-resolution microscopy (M. Boersma, (56)). Future studies will test this and related models to provide an understanding about the relationships among FtsZ treadmilling, PBP complex movement, and PG synthesis rate and location in different bacteria.

## Experimental Procedures

Detailed experimental procedures are described in *Supporting Information (SI) Experimental Procedures*, including bacterial strains and growth conditions; Western blotting; 2D-epifluorescence microscopy and demograph generation; growth and imaging of live cells by TIRFm; TIRFm image acquisition and processing; periodicity analysis of FtsZ filaments/bundles in nascent rings; culture growth and sample preparation for micro-hole immobilization of *Spn* cells; image acquisition, processing, and data analysis of vertically oriented cells in micro holes; single-molecule (SM) TIRFm; TIRFm of *ftsZ*(G107S)//P_Zn_-*ftsZ*-*sfgfp* merodiploid strains; TIRFm of HT-bPBP2x in an *ftsZ*(G107S) mutant; TIRFm of P_Zn_-*ftsZ*(D214A) or P_Zn_-*ftsZ*(D214A)-*sfgfp* merodiploid strains; 3D-SIM immunofluorescence microscopy (IFM); coimmunoprecipitation (co-IP) of FtsW-GFP with bPBP2x-FLAG; labelling of FtsZ(GTPase) mutant cells with FDAAs; and TIRFm of methicillin treated cells.

## Acknowledgements

We thank Jack Ryan and Jiaqi Zheng for strain constructions, members of the Winkler laboratory and Ethan Garner for critical discussions, Jim Powers for help with OMX microscopy, Luke Lavis for Fluor JF549, Michael VanNieuwenhze for TADA, Jan-Willem Veening and Yves Brun for strains and constructs, Orietta Massidda for anti-FtsZ antibody, and Dalia Denapite, Reinhold Bruckner, and Regine Hackenback for anti-PBP2x antibody. This work was supported by NIH grants R01GM113172, RO1GM114315, and RO1GM127715 (to MEW); NSF grant MCB1615907 (to SLS); Wellcome Trust and Royal Society Sir Henry Dale Fellowship grant 206670/Z/17/7 (to SH); NIH predoctoral Quantitative and Chemical Biology (QCB) training grant T32 GM109825 (to AJP); and NIH equipment grant S10RR028697 (to the Indiana University Bloomington Light Microscopy Center).

## Author Contributions

Conception or design of the study (AJP, YC, SLS, YT, MJB, SH, MEW); performed research (AJP, YC, YAY, SH, MEW); contributed reagents and tools (AJP, JBV, HCTT, CD), analyzed data (AJP, YC, SLS, MJB, SH, MEW); wrote manuscript (AJP, SLS, SH, MEW).

## References

1. Haeusser DP & Margolin W (2016) Splitsville: structural and functional insights into the dynamic bacterial Z ring. Nat Rev Microbiol 14(5):305–319.

2. den Blaauwen T, Hamoen LW, & Levin PA (2017) The divisome at 25: the road ahead. Curr Opin Microbiol 36:85–94.

3. Du S & Lutkenhaus J (2017) Assembly and activation of the *Escherichia coli* divisome. Mol Microbiol 105(2):177–187.

4. Coltharp C & Xiao J (2017) Beyond force generation: Why is a dynamic ring of FtsZ polymers essential for bacterial cytokinesis? BioEssays 39(1):1–11.

5. Meeske AJ, et al. (2016) SEDS proteins are a widespread family of bacterial cell wall polymerases. Nature 537(7622):634–638.

6. Sham LT, et al. (2014) Bacterial cell wall. MurJ is the flippase of lipid-linked precursors for peptidoglycan biogenesis. Science 345(6193):220–222.

7. Daniel RA, Harry EJ, & Errington J (2000) Role of penicillin-binding protein PBP 2B in assembly and functioning of the division machinery of Bacillus subtilis. Mol Microbiol 35(2):299–311.

8. Du S, Pichoff S, & Lutkenhaus J (2016) FtsEX acts on FtsA to regulate divisome assembly and activity. Proc Nat Acad Sci USA 113(34):E5052–5061.

9. Gamba P, Veening JW, Saunders NJ, Hamoen LW, & Daniel RA (2009) Two-step assembly dynamics of the *Bacillus subtilis* divisome. J Bacteriol 191(13):4186–4194.

10. Yao Q, et al. (2017) Short FtsZ filaments can drive asymmetric cell envelope constriction at the onset of bacterial cytokinesis. EMBO J 36(11):1577–1589.

11. Loose M & Mitchison TJ (2014) The bacterial cell division proteins FtsA and FtsZ self-organize into dynamic cytoskeletal patterns. Nat Cell Biol 16(1):38–46.

12. Bisson-Filho AW, et al. (2017) Treadmilling by FtsZ filaments drives peptidoglycan synthesis and bacterial cell division. Science 355(6326):739–743.

13. Yang X, et al. (2017) GTPase activity-coupled treadmilling of the bacterial tubulin FtsZ organizes septal cell wall synthesis. Science 355(6326):744–747.

14. Garner EC, et al. (2011) Coupled, circumferential motions of the cell wall synthesis machinery and MreB filaments in B. subtilis. Science 333(6039):222–225.

15. Lee TK, et al. (2014) A dynamically assembled cell wall synthesis machinery buffers cell growth. Proc Nat Acad Sci USA 111(12):4554–4559.

16. Monteiro JM, et al. (2018) Peptidoglycan synthesis drives an FtsZ-treadmilling-independent step of cytokinesis. Nature 554(7693):528–532.

17. Massidda O, Novakova L, & Vollmer W (2013) From models to pathogens: how much have we learned about *Streptococcus pneumoniae* cell division? Environ Microbiol 15(12):3133–3157.

18. Garcia PS, Simorre JP, Brochier-Armanet C, & Grangeasse C (2016) Cell division of *Streptococcus pneumoniae*: think positive! Curr Opin Microbiol 34:18–23.

19. Morlot C, Zapun A, Dideberg O, & Vernet T (2003) Growth and division of *Streptococcus pneumoniae*: localization of the high molecular weight penicillin-binding proteins during the cell cycle. Mol Microbiol 50(3):845–855.

20. Mura A, et al. (2017) Roles of the essential protein FtsA in cell growth and division in Streptococcus pneumoniae. J Bacteriol 199(3), e00608–16.

21. Morlot C, Noirclerc-Savoye M, Zapun A, Dideberg O, & Vernet T (2004) The D, Dcarboxypeptidase PBP3 organizes the division process of *Streptococcus pneumoniae*. Mol Microbiol 51(6):1641–1648.

22. Land AD, et al. (2013) Requirement of essential Pbp2x and GpsB for septal ring closure in *Streptococcus pneumoniae* D39. Mol Microbiol 90(5):939–955.

23. van Raaphorst R, Kjos M, & Veening JW (2017) Chromosome segregation drives division site selection in Streptococcus pneumoniae. Proc Nat Acad Sci USA 114(29):E5959-E5968.

24. Jacq M, et al. (2015) Remodeling of the Z-Ring Nanostructure during the *Streptococcus pneumoniae* Cell Cycle Revealed by Photoactivated Localization Microscopy. mBio 6(4) pii: e01108–15.

25. Tsui HC, et al. (2014) Pbp2x localizes separately from Pbp2b and other peptidoglycan synthesis proteins during later stages of cell division of *Streptococcus pneumoniae* D39. Mol Microbiol 94(1):21–40.

26. Fleurie A, et al. (2014) MapZ marks the division sites and positions FtsZ rings in Streptococcus pneumoniae. Nature 516(7530):259–262.

27. Holeckova N, et al. (2015) LocZ is a new cell division protein involved in proper septum placement in Streptococcus pneumoniae. mBio 6(1):e01700–01714.

28. Land AD, Luo Q, & Levin PA (2014) Functional domain analysis of the cell division inhibitor EzrA. PloS One 9(7):e102616.

29. Fadda D, et al. (2007) *Streptococcus pneumoniae* DivIVA: localization and interactions in a MinCD-free context. J Bacteriol 189(4):1288–1298.

30. Tsui HT, et al. (2016) Suppression of a Deletion Mutation in the Gene Encoding Essential PBP2b Reveals a New Lytic Transglycosylase Involved in Peripheral Peptidoglycan Synthesis in *Streptococcus pneumoniae* D39. Mol Microbiol 100(6), 1039–1065.

31. Rued BE, et al. (2017) Suppression and synthetic-lethal genetic relationships of Δ*gpsB* mutations indicate that GpsB mediates protein phosphorylation and penicillin-binding protein interactions in *Streptococcus pneumoniae* D39. Mol Microbiol 103(6):931–957.

32. Beilharz K, et al. (2012) Control of cell division in *Streptococcus pneumoniae* by the conserved Ser/Thr protein kinase StkP. Proc Nat Acad Sci USA 109(15):E905–913.

33. Li Y, et al. (2018) MapZ Forms a Stable Ring Structure That Acts As a Nanotrack for FtsZ Treadmilling in *Streptococcus mutans*. ACS Nano 12(6):6137–6146.

34. Barendt SM, et al. (2009) Influences of capsule on cell shape and chain formation of wild-type and pcsB mutants of serotype 2 *Streptococcus pneumoniae*. J Bacteriology 191(9):3024–3040.

35. Ducret A, Quardokus EM, & Brun YV (2016) MicrobeJ, a tool for high throughput bacterial cell detection and quantitative analysis. Nat Microbiol 1(7):16077.

36. Moreno-Gamez S, et al. (2017) Quorum sensing integrates environmental cues, cell density and cell history to control bacterial competence. Nat Commun 8(1):854.

37. Wilson M (2004) Microbial Inhabitants of Humans: Their Ecology and Role in Health and Disease (Cambridge University Press, Cambridge).

38. Thanedar S & Margolin W (2004) FtsZ exhibits rapid movement and oscillation waves in helix-like patterns in *Escherichia coli*. Curr Biol 14(13):1167–1173.

39. Holden S (2018) Probing the mechanistic principles of bacterial cell division with super-resolution microscopy. Curr Opin Microbiol 43:84–91.

40. Salvarelli E, et al. (2015) The Cell Division Protein FtsZ from *Streptococcus pneumoniae* Exhibits a GTPase Activity Delay. J Biol Chem 290(41):25081–25089.

41. Redick SD, Stricker J, Briscoe G, & Erickson HP (2005) Mutants of FtsZ targeting the protofilament interface: effects on cell division and GTPase activity. J Bacteriol 187(8):2727–2736.

42. Boersma MJ, et al. (2015) Minimal Peptidoglycan (PG) Turnover in Wild-Type and PG Hydrolase and Cell Division Mutants of Streptococcus pneumoniae D39 Growing Planktonically and in Host-Relevant Biofilms. J Bacteriology 197(21):3472–3485.

43. Taguchi A, et al. (2018) FtsW is a peptidoglycan polymerase that is activated by its cognate penicillin-binding protein. bioRxiv 10.1101/358663.

44. Du W, et al. (2000) Two active forms of UDP-N-acetylglucosamine enolpyruvyl transferase in gram-positive bacteria. J Bacteriol 182(15):4146–4152.

45. Slager J, Aprianto R, & Veening JW (2018) Deep genome annotation of the opportunistic human pathogen *Streptococcus pneumoniae* D39. Nucleic Acids Res 10.1093/nar/gky725.

46. Rowlett VW & Margolin W (2015) The Min system and other nucleoid-independent regulators of Z ring positioning. Front Microbiol 6:478.

47. Wu LJ & Errington J (2012) Nucleoid occlusion and bacterial cell division. Nat Rev Microbiol 10(1):8–12.

48. Kjos M & Veening JW (2014) Tracking of chromosome dynamics in live *Streptococcus pneumoniae* reveals that transcription promotes chromosome segregation. Mol Microbiol 91(6):1088–1105.

49. Manuse S, et al. (2016) Structure-function analysis of the extracellular domain of the pneumococcal cell division site positioning protein MapZ. Nat Commun 7:12071.

50. Lara B, et al. (2005) Cell division in cocci: localization and properties of the *Streptococcus pneumoniae* FtsA protein. Mol Microbiol 55(3):699–711.

51. Wheeler R, Mesnage S, Boneca IG, Hobbs JK, & Foster SJ (2011) Super-resolution microscopy reveals cell wall dynamics and peptidoglycan architecture in ovococcal bacteria. Mol Microbiol 82(5):1096–1109.

52. Coltharp C, Buss J, Plumer TM, & Xiao J (2016) Defining the rate-limiting processes of bacterial cytokinesis. Proc Nat Acad Sci USA 113(8):E1044–1053.

53. Harris LK & Theriot JA (2016) Relative Rates of Surface and Volume Synthesis Set Bacterial Cell Size. Cell 165(6):1479–1492.

54. Sperber AM & Herman JK (2017) Metabolism Shapes the Cell. J bacteriol 199(11).

55. Hussain S, et al. (2018) MreB filaments align along greatest principal membrane curvature to orient cell wall synthesis. eLife 7:e32471.

56. Soderstrom B, Chan H, Shilling PJ, Skoglund U, & Daley DO (2018) Spatial separation of FtsZ and FtsN during cell division. Mol Microbiol 107(3):387–401.

